# Sema3A Facilitates a Retrograde Death Signal via CRMP4-Dynein Complex Formation in ALS Motor Axons

**DOI:** 10.1101/774737

**Authors:** Roy Maimon, Lior Ankol, Romana Weissova, Elizabeth Tank, Tal Gradus Pery, Yarden Opatowsky, Sami Barmada, Martin Balastik, Eran Perlson

## Abstract

Amyotrophic Lateral Sclerosis (ALS) is a fatal neurodegenerative disease with selective dysfunction; it causes the death of motor neurons (MNs). In spite of some progress, currently no effective treatment is available for ALS. Before such treatment can be developed, a more thorough understanding of ALS pathogenesis is required. Recently, we demonstrated that ALS-mutated muscles contribute to ALS pathology via secretion of destabilizing factors such as Sema3A; these factors trigger axon degeneration and Neuromuscular Junction (NMJ) disruption. Here, we focus on the molecular mechanism by which muscle contribute to MNs loss in ALS. We identified CRMP4 as part of a retrograde death signal generated in response to muscle-secreted Sema3A, in ALS-diseased MNs. Exposing distal axons to Sema3A induces CRMP4-dynein complex formation and MN loss in both mouse (SOD1^G93A^) and human-derived (C9orf72) ALS models. Introducing peptides that interfere with CRMP4-dynein interaction in MN axons profoundly reduces Sema3A-dependent MN loss. Thus, we discovered a novel retrograde death signal mechanism underlying MN loss in ALS.

**Summary:** Maimon et al. identify a novel retrograde death mechanism that contribute to MN loss in ALS, in which CRMP4-Dynein complex is form and retrogradely move along the axon.

## Background

Amyotrophic lateral sclerosis (ALS) is a lethal neurodegenerative disease that affects upper and lower motor neuron (MN) health. This process leads to spasticity, muscle atrophy, and paralysis, which develop into respiratory failure and patient death (Frey *et al*., 2000; Fischer *et al*., 2004; Boillée, Vande Velde and Cleveland, 2006; Moloney, de Winter and Verhaagen, 2014; Peters, Ghasemi and Brown, 2015). Currently, no effective treatment exists for ALS, and finding basic mechanisms underlying the progressive cell death is crucial. Although the cause of most ALS cases is unknown (sporadic ALS-sALS), about 10% of ALS cases are inherited (familial ALS –fALS). The most common mutations responsible for fALS include expansions of a repeated DNA element (GGGGCC) in a gene termed *C9orf72*, and point mutations in the superoxide dismutase 1 (SOD1) gene (Rosen *et al*., 1993; DeJesus-Hernandez *et al*., 2011; Renton *et al*., 2011). The SOD1^G93A^ transgenic exhibit an ALS-like phenotype, including axon degeneration, cytoplasmic inclusions, gliosis, loss of body weight, as well as fatal upper and lower motor neuron deficits (Fischer *et al*., 2004). However, therapeutic success in the SOD1^G93A^ mouse model, as well as other ALS mouse models has not translated so far into an effective therapy. Thus, there is an urgent need to use other more effective models. The use of human induced pluripotent stem cells (iPSCs) from sporadic and familial ALS patients is a genetically precise and promising approach for elucidating disease mechanisms and testing therapeutic strategies. Indeed, in the past few years, multiple groups worldwide have characterized iPSC-derived MNs that have recapitulated many aspects of ALS pathology (Wen *et al*., 2014; Alves *et al*., 2015; Devlin *et al*., 2015; Lee and Huang, 2017; Fujimori *et al*., 2018; Shi *et al*., 2018).

A hallmark finding in ALS models and humans with ALS is alterations in the axonal transport process (Bilsland *et al*., 2010; Perlson *et al*., 2010; Gershoni-Emek *et al*., 2015; De Vos and Hafezparast, 2017). MNs are highly polarized cells with long axons that can reach 1 meter long in adult humans. In order to survive and function, MNs depend on proper delivery of information and signalling events along the axons from synapse to their soma (Harrington and Ginty, 2013; Millecamps and Julien, 2013; Terenzio, Schiavo and Fainzilber, 2017; Zahavi, Maimon and Perlson, 2017). The dynein/dynactin and kinesin motor protein families are responsible for retrograde and anterograde axonal transport, respectively (Paschal and Vallee, 1987; Howard, Hudspeth and Vale, 1989). Importantly, mutations in kinesin and dynein/dynactin are associated with ALS in humans (LaMonte *et al*., 2002; Münch *et al*., 2004; Steinberg *et al*., 2015; Nicolas *et al*., 2018). Several studies suggest that alterations in the cross talk and long-distance signaling pathways between neurons and their diverse extracellular cues, which are mediated by an axonal transport process, contribute to ALS pathology (Boillée, Vande Velde and Cleveland, 2006; Perlson *et al*., 2009; Gibbs *et al*., 2018).

Sema3A is a secreted protein that was initially identified because of its ability to induce the collapse and paralysis of axonal growth cones of chick sensory neurons *in vitro* (Luo, Raible and Raper, 1993). However, Sema3A has also been shown to induce neuronal cell death via several mechanisms (Nakamura, Kalb and Strittmatter, 2000; de Wit *et al*., 2006; Ben-Zvi *et al*., 2008; Yamashita *et al*., 2014; Wehner *et al*., 2016) and to be involved in a variety of pathologies in adults, including ALS (Good *et al*., 2004; De Winter *et al*., 2006; Venkova *et al*., 2014; Maimon *et al*., 2018). Collapsin Response Mediator Proteins (CRMPs) constitute a family well known for their intracellular response to Sema3A-Plexin binding. There are 5 known CRMPs in vertebrates, all of which share ∼75 percent sequence similarities (Schmidt and Strittmatter, 2007). The canonical signaling events of CRMPs during Semaphorin signaling involve their phosphorylation via GTPase activity, which further leads to microtubule destabilization and axon retraction (Sasaki *et al*., 2002; Yamashita and Goshima, 2012; Balastik *et al*., 2015). In addition to their role in mediating Sema3A intrinsic responses, CRMPs were also reported to bind dynein and kinesin, and modulate their function (Arimura *et al*., 2009; Rahajeng *et al*., 2010). Several studies further demonstrated the involvement of CRMPs in neurodegenerative diseases (Charrier *et al*., 2003; Yamashita and Goshima, 2012; Nagai, Baba and Ohshima, 2017). Specifically, CRMP4 expression levels were previously found to be elevated in the SOD1^G93A^ mouse spinal cord, and were suggested to hamper MN health (Duplan *et al*., 2010; Valdez *et al*., 2012; Nagai *et al*., 2015). However, the mechanism underline the involvement of CRMP4 mediating cell death and its role in other ALS model is still elusive. Moreover, mutations in CRMP4 were associated with ALS patients (Blasco *et al*., 2013).

Here, we identified a novel retrograde death signal, triggered by Sema3A, which facilitates MN loss in both murine and human ALS models. This process is mediated by the formation of CRMP4-Dynein complex along diseased axons. Interfering with the binding of CRMP4 to Dynein by using short blocking peptides rescues Sema3A-dependent MN loss in these ALS models.

## Results

### SOD^G93A^ MNs and muscle co-cultures enhance MN loss

MN cell death is a key process in ALS pathology. Taking into consideration our previous findings that muscle-dependent secretion of toxic factors can trigger axon degeneration in MNs and accelerate SOD1^G93A^ MN loss (Maimon *et al*., 2018), we investigated whether muscle toxicity could also hamper MN cell survival. Similarly as described before (Maimon *et al*., 2018; Ionescu *et al*., 2019), WT or SOD1^G93A^ P60 gastrocnemius muscles were dissected and cultured in the distal compartment of the microfluidic chamber for 7 days. Then, WT^ChAT::tdTomato^ or SOD1^G93A/ChAT::tdTomato^ explants were plated in the proximal compartment and grown in the Micro Fluidic Chambers (MFC) until axons traversed into the distal compartment and formed contact with the muscles. We tracked the number of ChAT-positive MNs within spinal cord explants at this point (10DIV), and 5 days after (15DIV) (Figure 1A-B). Our observations indicated that while under WT^ChAT::tdTomato^ conditions MNs survived, significant neuronal loss was observed when SOD1^G93A/ChAT::tdTomato^ explants were plated. Importantly, the MN loss was enhanced in the presence of SOD1^G93A^ muscles (Figure 1C) (Mean: WT-WT 91.85 ± 3.673; SOD1^G93A^-SOD1^G93A^ 69.24 ± 4.237; WT-SOD1^G93A^ 76.55 ± 2.575; SOD1^G93A^-WT 92.63 ± 3.047). Thus, although SOD1^G93A/ChAT::tdTomato^ MNs are prone to die, SOD1^G93A^ muscles also contribute to a more robust MN loss, suggesting that both intrinsic and extrinsic mechanistic defects contribute to ALS.

**Figure 1.**
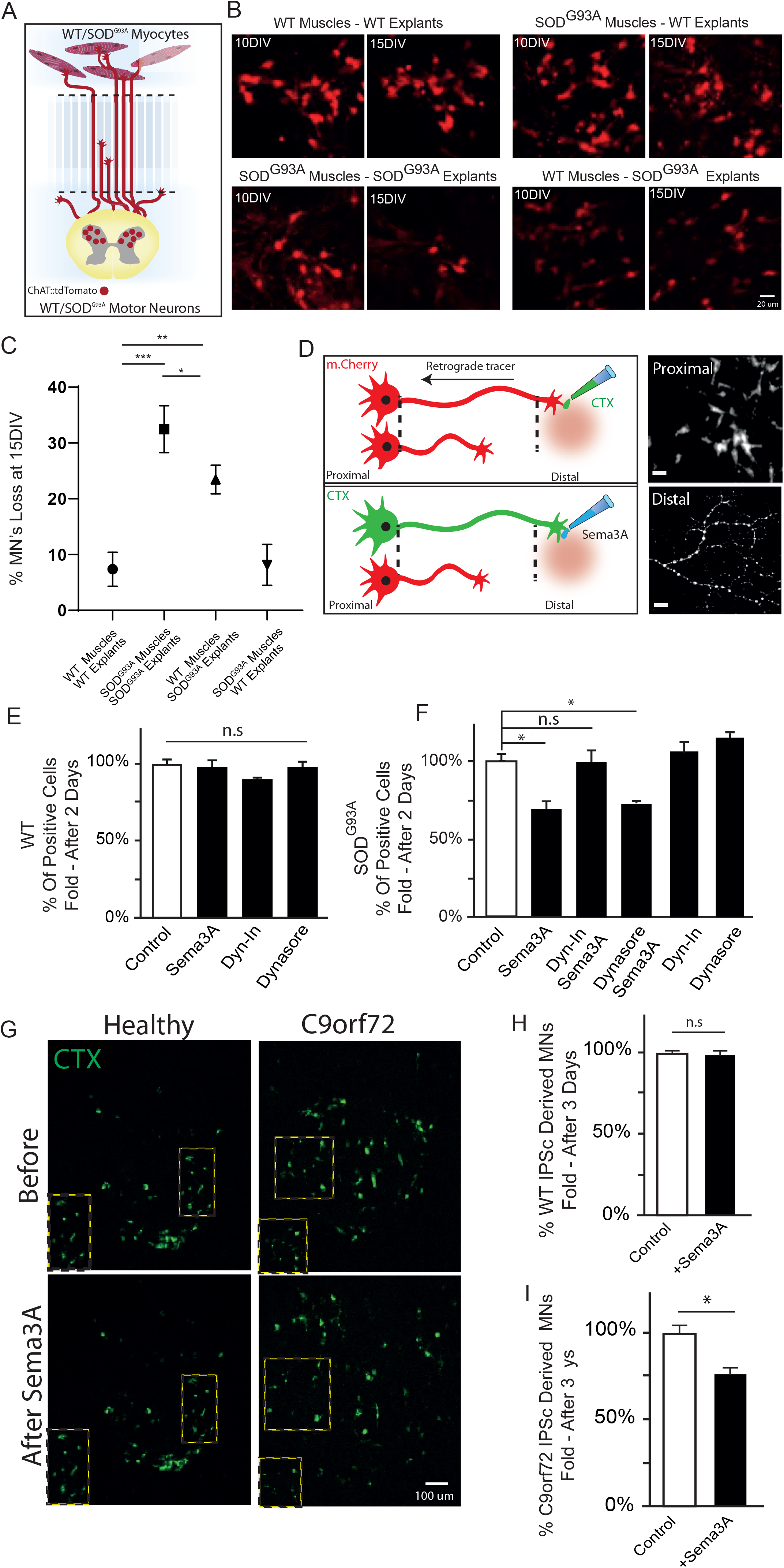
Sema3A activates a retrograde death signal in ALS mutated motor neurons. (A) Simplified illustration of the compartmentalized MFC used to culture WT^ChAT::tdTomato^ / SOD1^G93A/ChAT::tdTomato^ spinal cord explants together with WT/ SOD1^G93A^ primary myocytes in two different compartments. (B) Representative images of either SOD1^G93A/ChAT::tdTomato^ or WT^ChAT::tdTomato^ spinal cord explants in the proximal compartment of MFCs in four different co-culture combinations at 10DIV and at 15DIV. Images display significant motor neuron loss in SOD1^G93A/ChAT::tdTomato^ explant-cultured SOD1^G93A^ myocytes. Scale bar: 20 µm. (C) Quantification of the percentage of surviving cells per explant at 15DIV in co-culture compared to 10DIV (n = 3, One-way ANOVA, Tukey’s multiple comparisons test; p =0.0013, p=0.0001, p=0.036; WT-WT vs SOD1^G93A^-SOD1^G93A^, WT-WT vs WT-SOD1^G93A^, SOD1^G93A^-SOD1^G93A^ vs WT-SOD1^G93A^). (D) Left panel - Illustration of the experimental procedure for MNs in an MFC treated with the fluorescently tagged retrograde tracer CTX in the distal compartment. Neuronal cell bodies in the explant whose axons have traversed into the distal compartment were also labeled by the retrograde tracer. Right Panel - Representative images show CTX^+^ axons in the distal compartment and their cell bodies in the proximal compartment after applying Sema3A to the distal compartment. Gray denotes CTX tagging. Scale bar: 20µm. (E) Quantification of the CTX^+^ cell bodies in a WT explant 2 days after applying Sema3A to the distal compartment revealed no change in the cell count compared with the control. (F) Quantification of CTX^+^ in a SOD1^G93A^ explant 2 days after applying Sema3A to the distal compartment revealed a decreased number of CTX^+^ cell bodies, which can be blocked by a parallel application of the dynein inhibitor Cilliobrevin-D (Dyn-In) with Sema3A; however, a parallel application of Dynasore with Sema3A did not have a similar effect. Quantification is the percentage of total CTX^+^ cells after applying Sema3A compared to control medium; plots show mean ±SEM, (One-way ANOVA, Tukey’s multiple comparisons test, n = 3; *p<0.05, **p<0.01) dynein inhibitor and Dynasore treatments were used as a negative control. (G) Representative images of WT and C9orf72 Human IPSc-derived MNs before and after applying Sema3A, tagged with CTX using the same described technique. Green denotes CTX-positive cells. Scale bar: 100 µm. (H-I) Quantification of CTX^+^ in WT IPSc-derived MNs revealed no change in cell count 3 days after applying Sema3A distal compared with the control. However, applying Sema3A to the distal compartment of *C9orf72* iPSC-derived MNs resulted in a significant loss after 3 days. Quantification is the percentage of total CTX^+^ cells after applying Sema3A compared to control medium and is shown as the mean ±SEM (Student’s t-test, n=3, *p<0.05)

### Sema3A activates a retrograde death signal in MNs of SOD^G93A^ mice

As our previous report indicated, Sema3A is one of the toxic factors that are secreted in excess by ALS-diseased muscles. We also found that Sema3A facilitates axon degeneration and NMJ disruption (Maimon *et al*., 2018). Sema3A were shown to be involved in ALS also by other groups (De Winter *et al*., 2006; Venkova *et al*., 2014). Here, we hypothesized that Sema3A will lead not only to MN axon degeneration, but also to MN loss. To better understand this process, explants from WT embryos were cultured and grown in the MFC until axons traversed into the distal compartment. At this point, the retrograde tracer Alexa-647-conjugated Cholera Toxin Subunit B (CTX) was added to the distal compartment in order to exclusively label the cell bodies of neurons in the explant whose axons traversed into the distal compartment (Figure 1D). The number of CTX**^+^** cells in the explant was quantified before, and then daily, for one week, after Sema3A or its control medium was applied to the distal compartment (Figure 1E). Our observations of WT explants did not indicate any significant decrease in the number of cells (Figure 1E) (mean fold change over control: Sema3A 0.95 ± 0.06; control 1 ± 0.05). However, applying Sema3A similarly to explants from SOD1^G93A^ embryos resulted in a ∼30% reduction of CTX^+^ cells as early as after two days, compared with ∼5% in the control (Figure 1F)(mean fold change over control: Sema3A 0.68 ± 0.06; control 1 ± 0.04). Thus, despite inducing the degeneration of both healthy and diseased MN axons, Sema3A affects survival only in SOD1^G93A^ MNs.

Axonal transport is a process whereby cargo and signaling molecules are moved efficiently to and from the MN peripheral axon; it is performed in the retrograde direction by the dynein motor complex (Paschal and Vallee, 1987; Ibáñez, 2007; Zahavi, Maimon and Perlson, 2017). Previous studies have also shown that Sema3A can be internalized and undergoes retrograde transport to the soma (Dang, Smythe and Furley, 2012; Yamashita *et al*., 2014; Wehner *et al*., 2016). It has also been shown that Sema3A facilitates antero- and retrograde axoplasmic transport of organelles (Goshima *et al*., 1997). In order to determine whether Sema3A triggers MN loss in ALS via a retrograde signaling event, we used an inhibitor for Dynein motor protein (Ciliobrevin-D; hereafter Dyn-In) and thus blocked all retrograde transport events (Movie 1-2). We repeated our experiments in the MFC with SOD1^G93A^ explants, but this time Dyn-In was added to the distal compartment prior to Sema3A treatment (Fig 1F). Importantly, inhibiting retrograde transport in the distal axon prevented MN loss in the SOD1^G93A^ explant (mean fold change over control: Sema3A + Dyn-In 0.92 ± 0.09; control 1 ± 0.04), indicating that Sema3A propagates cell death via a retrograde signaling event. To further determine whether endocytosis of Sema3A components is important for death signaling, as was shown before in different neurons (Wehner et al., 2016), we applied Dynasore, a dynamin-dependent endocytosis inhibitor to the distal axons prior to applying Sema3A, which was shown to internalize via this pathway (Fournier *et al*., 2000; Castellani, Falk and Rougon, 2004)(Movie 3-4)(Supplementary Figure 1). Inhibiting Sema3A internalization did not rescue the MNs, and the number of CTX^+^ cells was significantly reduced by ∼30% after 2 days (Figure 1F)(mean fold change over control: Sema3A + Dynasore 0.71 ± 0.01; control 1 ± 0.04). Hence, these findings suggest that unlike in other types of neurons, Sema3A activates the retrograde transport of a secondary messenger in order to facilitate MN loss, which is independent of its internalization.

### C9orf72 patient-derived MN loss

Human iPSC-derived MN cultures are currently widely used as an *in vitro* model in studying ALS disease. Specifically, iPSC-derived MNs from individuals with *C9orf72* mutations display several disease-relevant phenotypes, including abnormal excitability and reduced survival (Devlin *et al*., 2015; Sances *et al*., 2016; Lee and Huang, 2017; Shi *et al*., 2018). We therefore plated human iPSC-derived MNs from healthy controls and C9orf72-ALS patients in MFCs (Supplementary Figure 2 A-C). Staining for the neuronal markers NFH and Tau, along with the MN-specific marker HB9, validated the character of our cultures (Supplementary Figure 2 D-F). Here too, we hypothesized that applying Sema3A to distal axons would trigger MN loss. To this end, we applied the retrograde tracer Alexa-647-conjugated Cholera Toxin Subunit B (CTX) after axons reached the distal compartment, labeling only the cell bodies of MNs whose axons traversed into the distal compartment. The number of CTX**^+^**iPSC-derived MNs was quantified before and 3 days after Sema3A was applied to the distal compartment (Figure 1 G). We did not detect any significant loss of WT iPSC-derived MNs in response to Sema3A, whereas *C9orf72* iPSC-derived MNs displayed a 25 percent cell loss upon treatment with Sema3A (Figure 1 G-H). Hence, these data suggest that Sema3A also triggers MN death in human iPSC-derived MNs carrying the *C9orf72* mutation.

**Figure 2.**
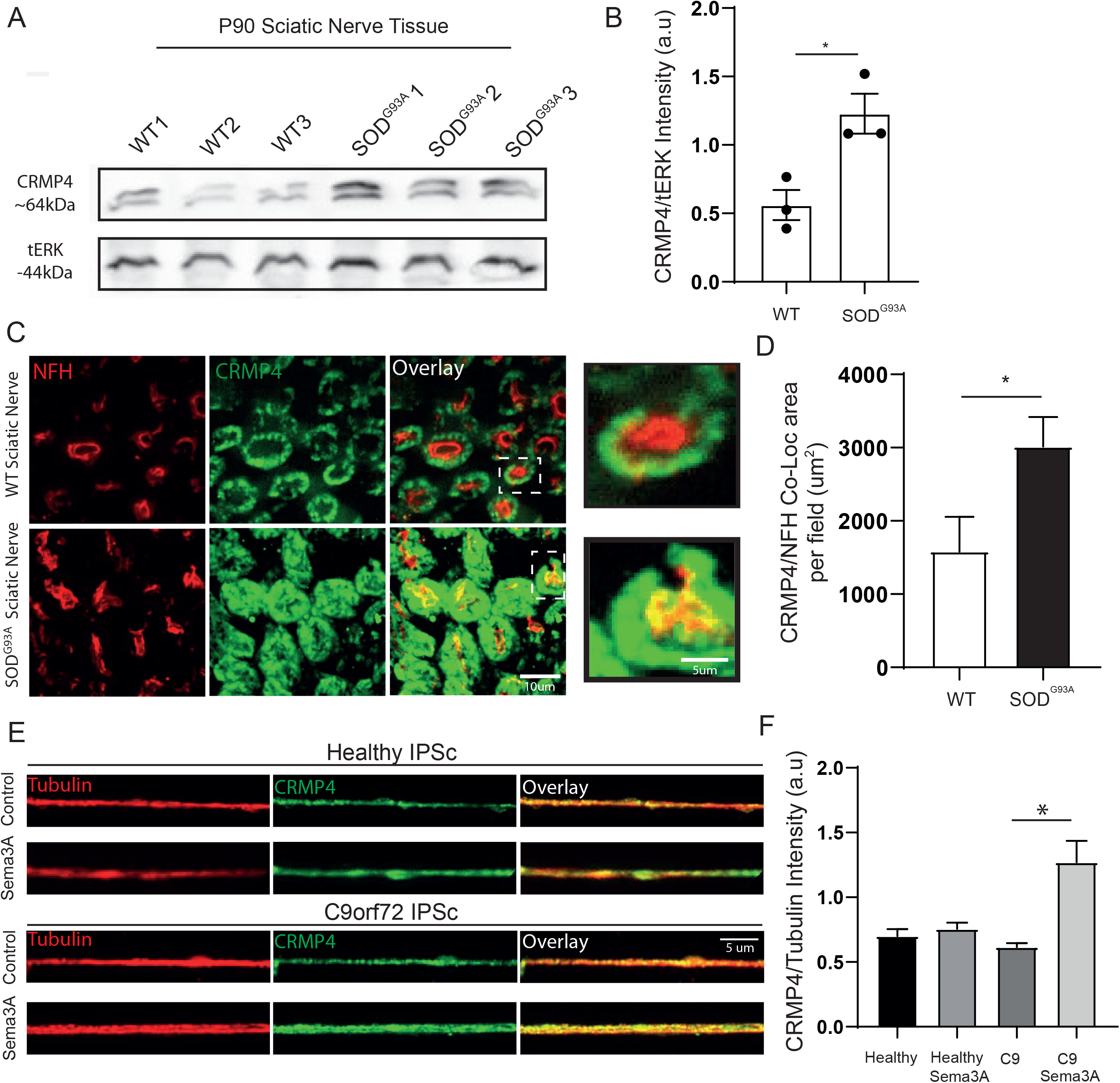
CRMP4 elevation in ALS models. (A-B) Western blot analysis and quantification of P90 SN tissues revealed that the levels of CRMP4 are elevated in *SOD1^G93A^*SN compared with their corresponding WT control. tERK was used as a loading control (Student’s t-test, n = 3, *p = 0.0215). (C) Representative images of fixed and immunostained SN cross sections showing elevation in the co-localization area of CRMP4 and NFH in the SOD1^G93A^ group compared with its WT. Red denotes NFH, Green denotes CRMP4. Left panel scale bar: 10 µm. Right panel is an inset - Scale bar: 5 µm. (D) Quantification of the images in C revealed a significant difference between the WT and *SOD1^G93^* conditions (Student’s t-test, n = 3, *p = 0.042). (E) Representative images of fixed and stained healthy or *C9orf72* iPSC-derived MNs treated with Sema3A or control in the distal compartment for 6 h. Red denotes Tubulin, Green denotes CRMP4. Scale bar: 5 µm. (F) Quantification of the images in E reveals a significant increase in the *C9orf72* iPSC-derived MN group treated with Sema3A. (One-way ANOVA, Tukey’s multiple comparisons test, n = 3, *p = 0.042.)

### CRMP4 levels are elevated in ALS models

Our data indicate that Sema3A facilitates a downstream signaling event that eventually forms a retrograde death signal in diseased MNs. Thus, we focused our study on well-established Sema3A downstream effectors, the CRMP protein family. The canonical signaling event by which CRMPs mediate the Sema3A repulsion effect is via microtubule destabilization. Interestingly, CRMP4 levels were recently found to be elevated in the SOD1^G93A^ spinal cord; this leads to MN cell death in unknown mechanism. Moreover, a rare mutation in CRMP4 was found in a population of ALS patients (Duplan *et al*., 2010; Valdez *et al*., 2012; Blasco *et al*., 2013; Nagai *et al*., 2015). Thus, we assumed that CRMP4 might be a key factor contributing to the Sema3A-dependent MN loss that we observed. In order to verify that levels of CRMP4 are indeed higher in SOD1^G93A^ sciatic nerve, we performed western blot analysis and compared WT and SOD1^G93A^ P90 sciatic nerve (SN) extracts (Figure 2A). We found a significant elevation in CRMP4 expression in SOD1^G93A^ SNs, compared with WT SNs (Figure 2B) (mean: WT 0.56 ± 0.11; SOD1^G93A^ 1.22 ± 0.14). We further immunostained P90 sciatic nerves and searched for CRMP4 alteration specifically in NF-H-positive axons (Figure 2C). We observed an increase in the number of positive NF-H and CRMP4 colocalized areas per field (mean in µm^2^: WT 1583 ± 478; SOD1^G93A^ 3015± 404) (Figure 2D). Our pervious data indicated that SOD1^G93A^ SNs are exposed to higher Sema3A expression (Maimon *et al*., 2018); therefore, we assumed that CRMP4 will also be elevated *in vitro* after Sema3A treatment in diseased MNs. We performed immunostaining in both healthy and diseased human iPSC-derived MNs, and assessed the change in CRMP4 levels in axons in the presence or absence of Sema3A. Our analysis revealed that indeed CRMP4 intensity was increased only in diseased MNs treated with Sema3A (mean: Healthy 0.7 ± 0.05; Healthy+Sema3A 0.75 ± 0.047; C9orf72 0.61 ± 0.03; C9orf72 1.27 ± 0.16) (Figure 2E-F). These results imply that intense Sema3A signaling may drive the rise in CRMP4 levels in diseased MNs, and specifically in axons.

### CRMP4-dynein complex formation in ALS models

In addition to mediating Sema3A signaling, CRMPs were previously reported to bind motor proteins and modulate their function (Arimura *et al*., 2009; Rahajeng *et al*., 2010). Hence, we further hypothesized that elevated CRMP4 levels mediate MN loss via its interaction with the motor protein dynein. We assumed that CRMP4-dynein cross talk would be enhanced in MN axons of ALS models that were exposed to Sema3A. To this end, we first extracted SN axoplasms from WT and SOD1^G93A^ P90 mice, and looked for CRMP4 that co-purified with dynein intermediate chain (DIC) via immunoprecipitation. Strikingly, we observed a strong interaction of DIC with CRMP4 exclusively in the SOD1^G93A^ mice (mean: WT 0.28 ± 0.11; SOD1^G93A^ 1.48 ± 0.62)(Figure 3A-B). We next used the Proximity Ligation Assay (PLA) to validate our result *in vitro*. As expected, we observed that indeed there is an increase in the CRMP4-dynein colocalization along axons of the Sema3A-treated group, indicating that Sema3A triggers this complex formation. Importantly, this colocalization was twice as high in the SOD1^G93A^-treated axons (Figure 3C-E). Next, we compared the co-localized CRMP4-dynein puncta in human iPSC-derived MNs and observed similar results. The CRMP4-dynein colocalization in *C9orf72* iPSC-derived MN axons after Sema3A treatment was significantly higher compared with healthy control iPSC-derived MN axons (mean puncta per axon: Healthy/control 5.4 ± 0.53; C9orf72/Sema3A 10 ± 1.2) (Figure 3 F-G). Hence, we demonstrated both *in vivo* and *in vitro* that CRMP4-dynein interaction increases in ALS-mutated MNs.

**Figure 3.**
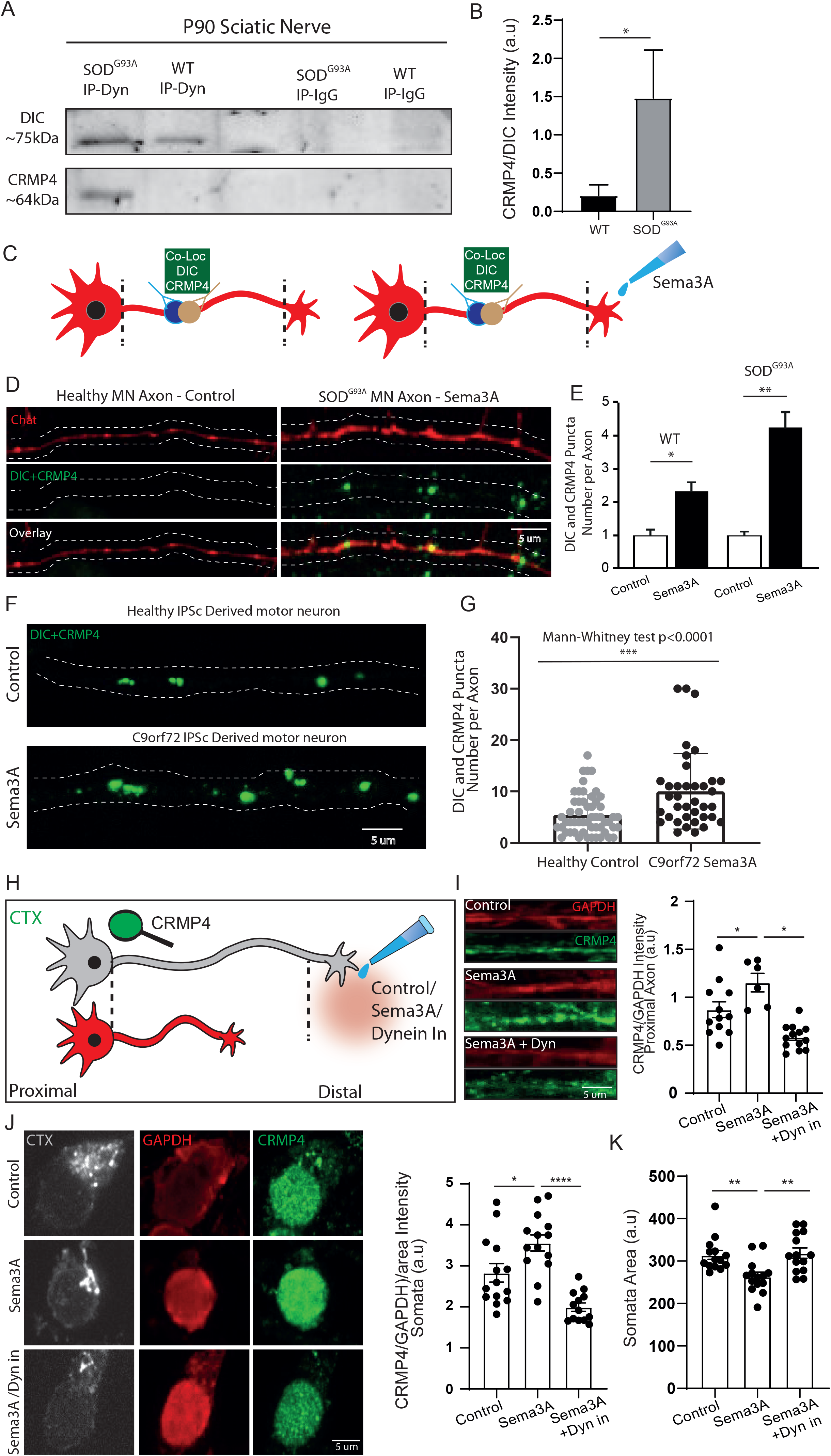
CRMP4 and dynein complex formation in ALS models. (A) Immunoprecipitation of DIC followed by western blot analysis of CRMP4 showing a strong association of DIC with CRMP4 in SOD1^G93A^ SNs compared to WT SNs under physiological conditions. IgG antibody was used as a control. (B) Quantification of 3 pull down repeats with anti-DIC revealed a significantly stronger interaction of DIC with CRMP4 in *SOD1^G93A^* compared with its WT. The CRMP4 intensity band was normalized to the DIC intensity band in each repeat. (Ratio Paired t-test, n = 3, 12 SN were used in each repeat, *p = 0.0416). (C) Illustration of the experimental procedure for the proximity ligation assay in an MFC. (D) Representative images of the Proximity ligation assay using CRMP4 and dynein antibodies in healthy and diseased mouse MNs that were exposed to either control or the Sema3A 8 h prior to fixation. (E) Quantification of the images in D revealed an increase in the CRMP4-DIC puncta number per axon in the Sema3A-treated group. CRMP4-DIC puncta number per axon was further increase when counting Sema3A-treated MNs carrying the *SOD1^G93A^* mutation (Student’s t-test, n = 3). (F) Representative images of the CRMP4-DIC puncta number per axon in human iPSC-derived MNs. (G) Bar graph shows an increase in the CRMP4-DIC puncta number in *C9orf72* iPSC-derived MNs that were treated with Sema3A compared with their healthy control (Mann Whitney test, n = 3, \**\*\**p = 0.0003). (H) Illustration of the experimental procedure for iPSC-derived MNs in an MFC treated with the fluorescently tagged retrograde tracer CTX in the distal compartment. Neuronal cell bodies whose axons have traversed into the distal compartment were marked by the retrograde tracer. Then, the distal axons were treated with Sema3A or control and the intensity of CRMP4 was measured either in the cell body or in the proximal axons. (I) Representative images (left panel) of proximal axons that were fixed and stained for CRMP4 and GAPDH after Sema3A/Sema3A dynein inhibitor or control treatment. Green denotes CRMP4, Red denotes GAPDH. Scale bar: 5um. (Right panel) Quantification of the images’ intensity revealed an increase in CRMP4 expression after Sema3A treatment compared with control treatment. This effect was diminished when dynein inhibitor was applied prior to Sema3A treatment, suggesting an increase in CRMP4 intensity due to retrograde movement. (J) Representative images (left panel) of cell somata that were fixed and stained for CRMP4 and GAPDH after Sema3A/Sema3A dynein inhibitor or control treatment. Gray denotes CTX, Green denotes CRMP4, Red denotes GAPDH. Scale bar: 5um. (Right panel) Quantification of the images’ intensity revealed an increase in CRMP4 expression after Sema3A treatment compared with control treatment that was reduced after dynein inhibitor application, (n = 3, One-way ANOVA, Tukey’s multiple comparisons test; *p =0.013, *p=0.0001). (K) Quantification of the cell somata area under the same conditions as in J shows that Sema3A treatment in the distal axons results in a decreased cell area. However, this effect was diminished when dynein inhibitors were applied prior to Sema3A treatment (n = 3, One-way ANOVA, Tukey’s multiple comparisons test; **p =0.0067, **p=0.0031).

Since dynein is a retrograde motor protein, we expected that the dynein interacting/cargo proteins would be retrogradely transported. We therefore assumed that CRMP4 will change its localization and will move along the axon towards the cell body. To better understand this process, we attempted to track exogenous CRMP4-GFP movement in live cells, but overexpressing CRMP4-GFP resulted in a uniformly defuse distribution. However, we hypothesized that endogenous CRMP4 would demonstrate retrograde transport that could be identified by immunostaining distinct cellular compartments (Olenick, Dominguez and Holzbaur, 2019). To this end, we specifically examined CRMP4 levels in the cell bodies and proximal axons of MNs that were distally exposed to Sema3A, Sema3A + Dyn-In, or control by using the same CTX tagging approach as described before (Figure 3H). Similarly to what we have previously shown, we observed that the levels of CRMP4 in proximal axons and cell bodies were elevated by Sema3A treatment. Strikingly, inhibiting dynein completely blocked the subsequent elevation of CRMP4 by Sema3A (mean: control 2± 0.2; Sema3A 3 ± 0.1; Sema3A/Dyn-In 1 ± 0.01) (Figure 3I-J). Furthermore, we quantified the cell body area for each treatment group and observed that Sema3A-treated MNs exhibit smaller soma areas compared with control MNs. Here again, when Dyn-In was added prior to applying Sema3A, this effect was abolished (mean: control 314± 10; Sema3A 263 ± 10; Sema3A and Dynein inhibitor 318 ± 12) (Figure 3K). These data indicate that the elevation in CRMP4 in the cell bodies and in proximal axons is indeed mediated by dynein binding to axonal CRMP4 and subsequent retrograde transport of the CRMP4-dynein complex.

### Specific peptides interfere with CRMP4-dynein interaction

CRMP2 members were previously found to bind dynein motor protein (Arimura *et al*., 2009). Arimura et al. also characterized two specific domains in CRMP2 protein, which are responsible for dynein binding (Arimura *et al*., 2009). Since CRMP2 and CRMP4 share most of their sequence, we hypothesized that the CRMP2 dynein-binding domains would play a similar role in CRMP4. Following this idea, and on the basis of the CRMP4 protein crystal structure (PDB code 4CNT) (Ponnusamy *et al*., 2014) (Figure 4A), we first examined if the same domains that are responsible for CRMP2 dynein binding function similarly in CRMP4. To this end, we overexpressed full length GFP-CRMP4 or CRMP4 missing amino acids 100-150 (GFP-ΔCRMP4) in COS7 cells, and immunoprecipitated endogenous DIC. Western blot analysis of these fractions revealed a strong interaction of DIC with full-length GFP-CRMP4 but not GFP-ΔCRMP4 (Figure 4B-C)(mean: GFP-CRMP4 1.473± 0.373; GFP-ΔCRMP4 0.009± 0.001). Because a large truncation can result in CRMP4 misfolding and dysfunction, we pursued a different strategy and blocked the association between CRMP4 and dynein using small peptides modeled after the interaction domain. We designed 4 peptides (CRMP4 Binding Peptides – CBP1-4) from within the 50 amino acid domain with the potential to block the CRMP4-dynein interaction. Importantly, the designed peptides’ sequences are positioned away from the CRMP4 canonical tetramer, and therefore are not likely to compromise activities related to the protein’s homomeric assembly (Figure 4A). Strikingly, pre-incubating a mixture of CBP1-4 with lysate from GFP-CRMP4 overexpressing COS7 cells significantly reduced the CRMP4-DIC interaction (Figure 4 B-C; mean: GFP-CRMP4 1.473± 0.373; GFP-CRMP4 + Peptides 0.3± 0.05). We further examined the peptides’ ability to block the CRMP4-Dynein interaction in COS7 cells overexpressing Flag-tagged CRMP4. After an overnight incubation of cell lysate with CBP1-4, we pulled-down Flag-CRMP4 and blotted for dynactin (p150). Here again, application of the peptide mixture resulted in a dramatic decrease in CRMP4 binding to dynein complex (mean: control 0.77± 0.1; All Pep 0.17± 0.06) (Supplementary Figure 3A-B). We then determined whether introducing individual peptides might be sufficient to block the CRMP4-Dynein interaction. Using the same technique as before, we found that only CBP-4 had a mild but significant ability to block CRMP4’s interaction with dynactin (mean: control 1.92± 0.2; CBP1 1.44 ± 0.3; CBP2 1.34 ± 0.3; CBP3 1.8 ± 0.8; CBP4 1.13 ± 0.14) (Supplementary Figure 3 C-D). Hence, although CBP-4 seems to be sufficient for blocking CRMP4-Dynein binding, for achieving the optimal blocking of CRMP4-dynein complex formation, a combination of all four peptides is required.

**Figure 4.**
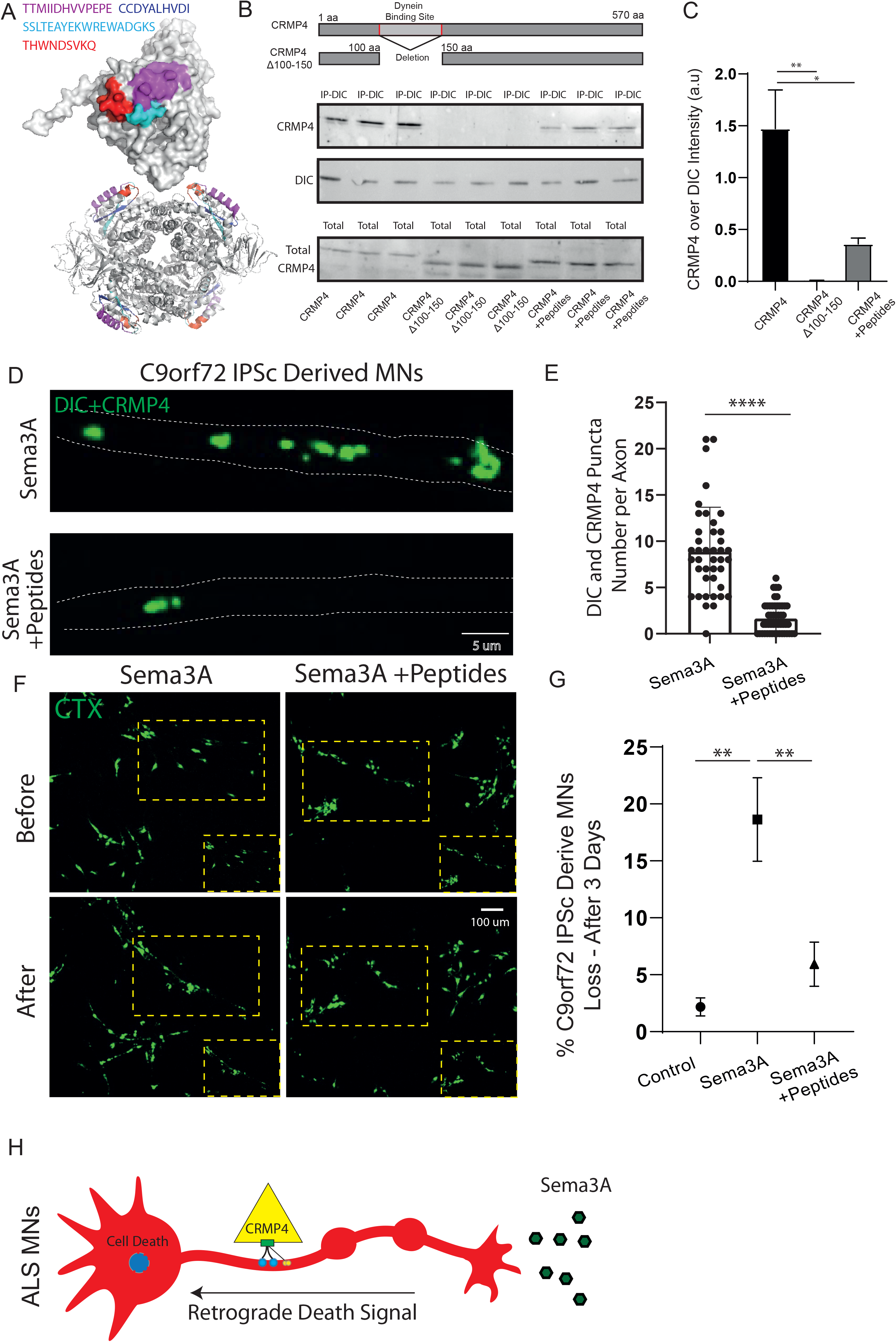
Specific peptides interfere with CRMP4-dynein interaction and rescue MN cell death. (A) Crystal structure (PDB code 4CNT) of CRMP4 monomer (upper panel) and biological tetramer assembly (lower panel). The derived peptides that were selected are highlited and color coded as indicated. Note that the peptides’ sequences are located on the surface of the protein away from the intermolecular tetramization interfaces. (B) Upper panel - immunoprecipitation of DIC followed by western blot analysis of CRMP4 showing a strong association of dynein complex with CRMP4 in COS7 cells overexpressing GFP-CRMP4. This binding completely abolished after deletion of amino acid 100-150 and can be interrupted with by pre-incubation of the protein extract with a 10 um mixture of CBP1-4. Lower panel - Total protein western blot analysis indicates similar protein levels before the pull-down assay. (C) Quantification of the blot in B shows a significant decrease in CRMP4-dynactin binding with GFP-ΔCRMP4 and with CBP1-4 pre-incubation, compared with control. The dynactin intensity band normalized to the Flag-CRMP4 intensity band in each repeat (One-way ANOVA, Tukey’s multiple comparisons test **p=0.007,*p=0.02, n=3). (D-E) Representative images and quantification of the proximity ligation assay for CRMP4-dynein with and without peptides reveal a decrease in the puncta number in C9orf72 iPSC-derived MNs after peptide application (Student’s t-test, p=0.0001, n=3). (F-G) Representative images and quantification of C9orf72 iPSC-derived MNs in the proximal compartment of an MFC before and after Sema3A treatment with and without 10 ug CBP1-4. The data indicate a significant 25% MN loss after applying Sema3A, which was prevented when Sema3A and the peptide mixture were introduced together. Green denotes CTX-positive cells. Scale bar: 100 um (n = 3, One-way ANOVA, Tukey’s multiple comparisons test; **p =0.004, **p=0.004). (H) Model - Sema3A mediates MN loss via CRMP4-Dynein complex formation. Blocking this interaction with specific peptides rescues human diseased iPSC-derived MNs from this fate.

Since CRMP4 retrograde transport occurs in distal axons of MNs, we decided to block CRMP4-dynein interaction specifically in MN distal axons. To this end, we plated human iPSC-derived MNs in MFCs and introduced CBP1-4 into the distal compartment comprised entirely of distal axons (Figure 4D). Using TAMRA peptides as a positive control, we observed that CBP1-4 peptides were successfully taken up by distal axons of MNs (Supplementary Figure 3 E-F). Strikingly, application of CBP1-4 significantly interfered with the interaction of CRMP4 and dynein in distal *C9orf72* iPSC-derived MN axons, but not control MN axons, as determined by PLA (mean: C9orf72 8.9± 0.7; C9orf72 with Pep 1.6± 0.2) (Figure 4E). In light of these data, we asked whether blocking the CRMP4-Dynein interaction in distal axons is sufficient to rescue ALS-diseased MN death in response to Sema3A. Introducing the peptides into diseased iPSC-MN axons prior to applying Sema3A completely abolishes Sema3A-dependent death (mean: Control 2.1± 0.7; Sema3A 18.6± 3.6; Sema3A + Pep 5.9± 1.9) (Figure 4F-G). Thus, CBP1-4 insertion ultimately into MN axon is sufficient to block CRMP4-Dyn interaction and eventually rescue Sema3A dependent MN loss

## Discussion

In this work we demonstrated that Sema3A mediates MN loss via CRMP4-dynein complex formation. Blocking this interaction with specific peptides rescued human diseased iPSC-derived MNs from this fate (Figure 4 H). However, open questions remain in this regard.

Our results, as well as those from other groups, suggest that Sema3A presence near the axons facilitate MN axon degeneration and cell death in ALS pathology (De Winter *et al*., 2006; Körner *et al*., 2016; Maimon and Perlson, 2019). Moreover, injecting specific antibodies against NRP1 or manipulating Sema3A expression using miR126-5p resulted in improvement in disease progression *in vivo* and *in vitro* (Venkova *et al*., 2014; Maimon *et al*., 2018). In contrast to these results, a recent study demonstrated that crossing mice expressing a truncated form of Sema3A with SOD1^G93A^ mice does not result in any rescue effect (Moloney *et al*., 2017). An explanation for this contradiction could be the possibility that secreted factors such as Sema3A play a more complex role in the biology of MNs (Zahavi, Maimon and Perlson, 2017). Indeed, Sema3A was shown to increase the survival of MNs when added to MN cultures, (Molofsky *et al*., 2014; Birger *et al*., 2018). Consistent with this, deletion of the Sema3A gene specifically in spinal astrocytes resulted in a gradual loss of spinal MNs (Molofsky *et al*., 2014). Apparently, Sema3A may play a dual spatial role in MNs: When introduced near MN axons at NMJs, it mediates their destabilization; however, when it is secreted by spinal astrocytes and targets MN soma, it acts as a survival factor. Thus, when targeting Sema3A for ALS, this aspect must take in consideration.

We further report that CRMP4 is elevated in ALS models. However, what mechanisms are responsible for CRMP4 elevation specifically in ALS is unknown. miRNA downregulation and defects in local protein synthesis are common features in several ALS models (Haramati *et al*., 2010; Costa and Willis, 2018). Along with that, Sema3A was shown to induce axonal local synthesis in several neuronal systems (Campbell and Holt, 2001; Wu *et al*., 2005; Manns *et al*., 2012; Cagnetta *et al*., 2018, 2019). Thus, we hypothesize that the elevations in CRMP4 that we observed are possibly due to increase in axonal protein synthesis. Specifically, our recent published work suggest that miR126-5p is downregulated in both muscles and MN axons of several ALS models (Rotem *et al*., 2017; Maimon *et al*., 2018). Thus, it is attempting to speculate further that miR126-5p downregulation mediates CRMP4 increases via local protein synthesis and consolidation of retrograde death signals in ALS models. However, further experiments should prove this idea.

Our data further suggest that CRMP4 forms complexes with dynein along ALS-diseased MNs and leads to their loss. However, it is still not clear whether CRMP4 itself activates the apoptotic program or whether it plays a regulatory role by mediating the arrival of the death complex. Since CRMP members were not yet reported to act as transcriptional factors, we assumed that CRMP4 is indeed only one protein in a bigger retrograde signaling complex. For example, JNK and c-Jun might also be part of this death signal mediated by Sema3A in ALS MNs, since it was previously shown that JNK signaling is elevated in ALS models and that it is a part of a retrograde death signal (Perlson *et al*., 2010; Escudero *et al*., 2019). Another possibility is that the neurotrophic receptor p75^NTR^ is also involved in this process. p75^NTR^ has been shown to regulate a diverse range of cellular functions including axon pruning (Singh *et al*., 2008) and neuronal death (Bamji *et al*., 1998; Kenchappa *et al*., 2010; Pathak *et al*., 2018). P75^NTR^ is retrogradely transported along the axon (Deinhardt *et al*., 2006; Harrington and Ginty, 2013; Cosker and Segal, 2014) and plays a role in generating a retrograde apoptotic signal that activates JNK (Kenchappa *et al*., 2010). It is noteworthy that the activity of Sema3A and its receptor was recently linked to p75^NTR^ (Ben-Zvi *et al*., 2007). Another possible candidate that was recently shown to be coupled with Sema3A is PTEN (Chadborn *et al*., 2006). Furthermore, it was established that the p75^NTR^-dependent apoptosis signal is promoted by PTEN activation (Song *et al*., 2010). Thus, future experiments should examine whether PTEN, p75^NTR^, and JNK indeed participate in Sema3A-dependent retrograde death signals in ALS.

Downregulation of CRMP4 has been previously suggested to improve neurodegeneration in *in vitro* and *in vivo* ALS models (Charrier *et al*., 2003; Duplan *et al*., 2010). Furthermore, *CRMP4* mutation has been associated with ALS in patients (Blasco *et al*., 2013), yet the mechanism of its action in ALS MNs has so far been elusive. Here, we demonstrate a specific mechanism by which Sema3A-induced CRMP4 binding to dynein is toxic in ALS. It would be interesting to determine whether disease-associated *CRMP4* mutations increase the interactions of CRMP4 with dynein. Indeed, the peptides we used are predicted to bind specifically at the site where CRMP4 mutation occurs (Blasco *et al*., 2013). Taken together, CRMP4 retrograde death-promoting signaling may be key to MN loss in ALS.

## Methods

### Animals

SOD1^G93A^ (Stock No. 002726) mice were originally obtained from Jackson Laboratories, and maintained by breeding with C57BL/6J mice. B6;129S6-Chat^tm2(cre)Lowl^/J (Stock No. 006410) and B6;129S6Gt(ROSA)26 Sor^tm14(CAG−tdTomato)Hze^/J (Stock No. 007908) mice were originally obtained from Jackson Laboratories. Animals were cross-bred in the Tel-Aviv SPF animal unit to yield homozygous ChAT::Rosa^tdTomato^ mice. The ChAT::Rosa^tdTomato^ colony was maintained by in-breeding males and females from the colony. The ChAT::Rosa^tdTomato^ colony was cross-bred with SOD1^G93A^ to yield SOD1^G93A/ChAT::tdTomato^ mice. C57BL/6 J mice were used as a WT mouse strain. Mice were genotyped using the PCR reaction (KAPA Bio Systems - Wilmington, MA, USA). DNA samples were generated from mouse ear or tail. Animal experiments were performed under the supervision and approval of the Tel-Aviv University Committee for Animal Ethics.

### IPSc Cultures

Healthy/Control iPSC lines, provided by Dr. Sami Barmada, were created and characterized as before (Tank *et al*., 2018). Two lines from fALS patients carrying the C9orf72 mutation, and two lines from healthy controls, were used for all experiments.

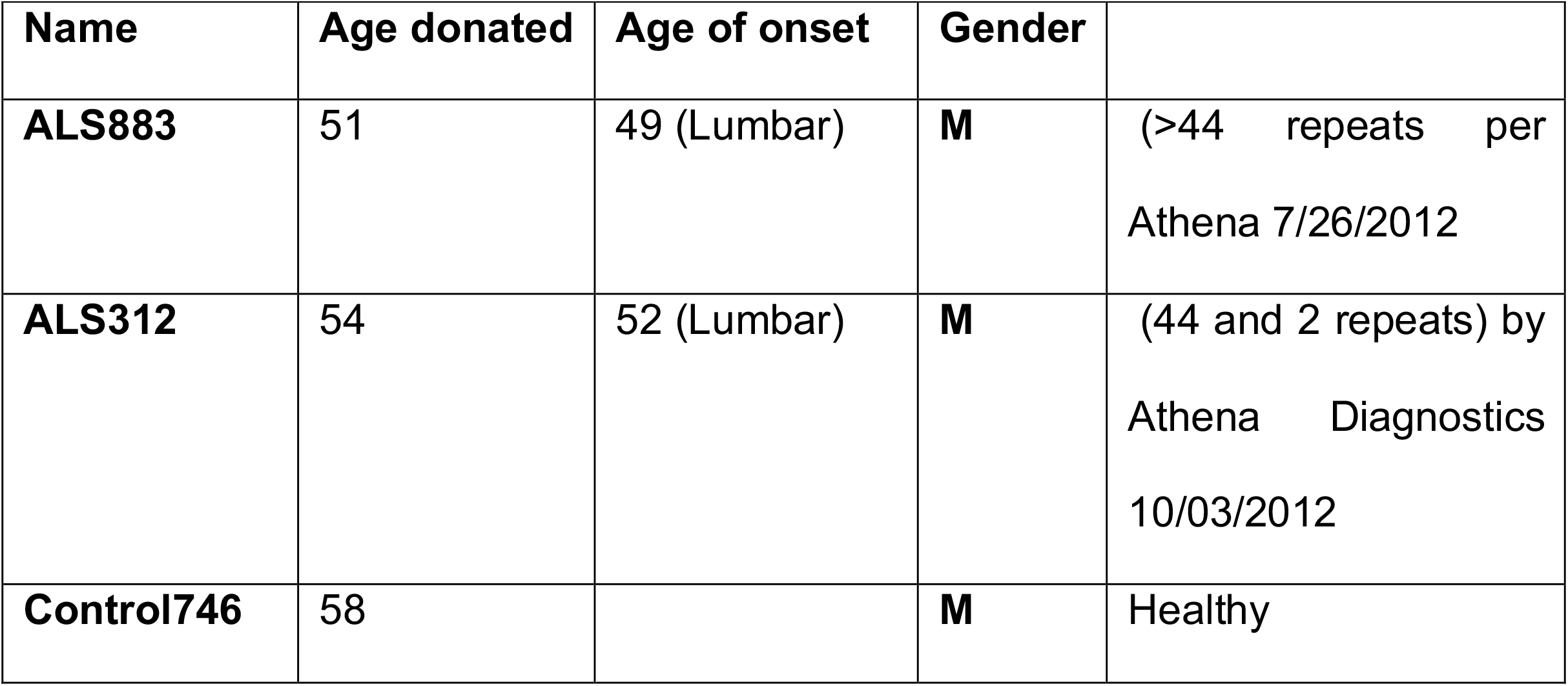
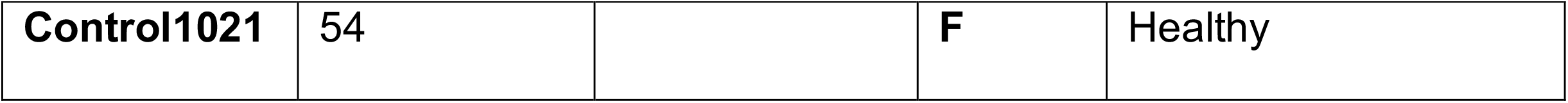

Colonies were groomed daily until each well of the 6-well plate was between 30% and 40% confluence and no spontaneously differentiated cells were observed. At this point, we used the direct iMN” (diMN) differentiation in monolayers from hiPSCs protocol published by Cedar Sinai and approved by Dhruv Sareen (Protocol number: CSMNC-SOP-C-005) for our experiments. Briefly, we induced MN differentiation by <u>MN Differentiation Stage 1 media</u>: prepared with – IMDM (LifeTech), F12 (LifeTech), NEAA (Gibco), B27 (LifeTech), N2 (LifeTech), PSA (LifeTech), LDN193189 0.2 µM (Selleck), SB431542 10 µM (Tocris), and CHIR99021 3 µM (Cayman Chemicals). The media was gently added to the wells, and colonies were grown with it for 5 days. <u>MN differentiation Stage 2</u>: At day 6 the colonies were dissociated, using accutase, and 100K cells were plated in the proximal compartment of our micro fluidic device. Stage 2 media, which contains IMDM, F12, NEAA, B27, N2, PSA, LDN193189 0.2 µM (Selleck), SB431542 10 µM (Tocris), and CHIR99021 3 µM (Cayman Chemicals), All-trans RA 0.1 µM (Stemgent), and SAG 1 µM (Sonic Hedgehog Agonist – Cayman Chemicals) media was added to both the distal and proximal compartments of the MFC and refreshed every 2 days until day 11. <u>MN differentiation Stage 3</u>: At day 12 the media was changed to stage 3 media prepared with IMDM, F12, NEAA, B27, N2, PSA, Compound E 0.1 µM (Calbiochem), DAPT 2.5 µM (Cayman Chemicals), db--cAMP 0.1 µM (Millipore), Alltrans RA 0.5 µM (Stemgent), SAG 0.1 µM, Ascorbic Acid 200 ng/ml (Sigma), BDNF 10 ng/ml (Alomone lab), and GDNF 10 ng/ml (Alomone lab) and was refreshed every 2 days until cells exhibited MN neuronal morphology and positive markers. Human iPSC experiments were performed under the supervision and approval of the Tel-Aviv University Committee for Human Ethics.

### Microfluidic chamber preparation

Polydimethylsilxane (PDMS) microfluidic chambers (MFCs) were designed and cast as described previously (Ionescu *et al*., 2016). Briefly, MFCs were fabricated from our designed templates and made from PDMS mixture at 70°C. After the wells were punched, a small ‘cave’ was made in the explant well near the grooves using a 25G needle, keeping the explant in place. Microfluidic devices were cleaned of surface particles using adhesive tape and were sterilized in 70% ethanol for 15 minutes. Devices were completely dried under sterile conditions using UV radiation, attached to a sterile 60-mm plastic dishes (Nunc) with gentle pressure and margins were sealed with PDMS before incubation at 60°C for 30 minutes to prevent the chamber from detaching. Muscle channels were coated with Matrigel diluted 1:10 with DMEM containing 2.5% PSN for 30 minutes at 37°C, before filling the muscle wells with 150µL of Bioamf-2 medium. The explant well and channel were filled with 150µL of 1.5 ng/mL polyornithine (P-8638, Sigma) in PBS overnight, and then replaced with 150 µL laminin (L-2020, Sigma), 1:333 in deionized distilled water (DDW) overnight. One day before plating the spinal cord explant, laminin was replaced with explant medium containing Neurobasal (Life Technologies) supplemented with 2% B27 (Invitrogen), 1% penicillin-streptomycin (Biological Industries), 1% Glutamax (Life Technologies), 25 ng/mL brain-derived neurotrophic factor (Alomone Labs), until the day on which co-culturing began.

### Motor neuron cell culture

Primary spinal cord neurons were cultured using E12.5 mouse embryos of either sex as previously described (Zahavi *et al*., 2015). Briefly, spinal cords were excised, trypsinized, and triturated. Supernatant was collected and centrifuged through a 4% BSA cushion. The pellet was resuspended and centrifuged through an Optiprep gradient (10.4% Optiprep (Sigma-Aldrich), 10 mM Tricine, 4% glucose) for 20 min at 760 x g with the brake turned off. Cells were collected from the interface, washed once in complete medium, and then plated in coated growth chambers. Cells were maintained in Complete Neurobasal Medium (Gibco) containing B27 (Gibco), 10% (v/v) horse serum (Biological Industries), 25 nM beta-mercaptoethanol, 1% Penicillin-Streptomycin (PS; Biological Industries), and 1% GlutaMAX (Gibco) supplemented with 1 ng/mL Glial-Derived Neurotrophic Factor (GDNF), 0.5 ng/mL Ciliary Neurotrophic Factor (CNTF), and 1 ng/mL Brain-Derived Neurotrophic Factor (BDNF), (Alomone Labs). Prior to plating, growth plates were coated with 1.5 g/mL poly D-L-ornithine (PLO; Sigma-Aldrich) overnight at 37 °C and with 3 g/mL Laminin (Sigma-Aldrich) for 2 hours at 37 °C. For immunofluorescence staining, 10,000 cells were plated on cover slides in 24-well plates. Cells were grown at 37 °C in 5% CO_2_.

### Spinal cord explants

Spinal cords were dissected from E12.5 mouse embryos of both sexes, either using HB9::GFP or SOD1^G93A^ stripped of meninges and dorsal root ganglia. The ventral horn was separated from the dorsal horn by longitudinal cuts along the spinal cord, and transverse sections up to 1 mm were placed in the explant well. Prior to plating, growth chambers were coated with 1.5 g/mL PLO overnight at 37 °C and 3 g/mL Laminin overnight at 37 °C. Explants were maintained in Spinal Cord Explant Medium containing Neurobasal, 2% B27, 1% PS, and 1% GlutaMAX, supplemented with 25 ng/mL BDNF. Explants were grown at 37 °C in 5% CO_2_.

### Primary myocyte culture

Skeletal muscle cultures were derived from the gastrocnemius muscle of adult P60 female mice of either SOD1^G93A^ background or their LM using techniques previously described (Ionescu *et al*., 2016). Briefly, gastrocnemius muscles were excised and incubated in 2 mg/mL collagenase I (Sigma-Aldrich) in DMEM containing 2.5% penicillin-streptomycin-nystatin (PSN, Biological Industries) for 3 hours. Muscles were then dissociated and incubated for 3 days in Matrigel-coated (BD Biosciences) six-well plates with Bioamf-2 medium (Biological Industries) with 1% PSN at a density of ∼120 myofibers per well. For purification of the myoblasts, adherent cells were trypsinized and pre-plated in an uncoated dish for 1 h at 37°C. Non-adherent cells were then transferred into a Matrigel-coated dish with Bioamf-2 medium. Pre-plating was repeated for two days, keeping the culture at less than 50% confluence, before plating cells in the MFC. Cultures were maintained at 37°C and 5% CO_2_. After the final pre-plating, 100,000 myocytes were cultured in the pre-coated distal compartment of the MFC.

### Fluorescence microscopy and image analysis

All confocal images were captured using a Nikon Ti microscope equipped with a Yokogawa CSU X-1 spinning disc and an Andor iXon897 EMCCD camera controlled by Andor IQ3 software. All live-imaging assays were performed in a humidified incubation chamber at 37°C, 5% CO_2_. Images were analyzed using ImageJ software.

### Rertrograde labeling of cell bodies in the MFC

Alexa Fluor 647-conjugated Cholera toxin subunit B (CTX; Thermo-Fisher C-347777) at 500 ng/mL was applied to the distal compartment of an MFC system while maintaining a higher liquid volume in the proximal compartment to prevent unspecific labeling by diffusion. After 8 hours, only somata whose axons traversed to the distal compartment were labeled.

### Recombinant Sema3A application

Recombinant Sema3A (R&D,1250-S3-025) at 500 ng/mL was used in our experiments. We dilute the Sema3A in poor neurobasal (PNB) containing Neurobasal medium (Gibco) with 1% PS and 1% Glutamax.

### Retrograde transport inhibition

In order to inhibit dynein-dependent retrograde transport, Cilliobrevin-D (Merck-Millipore, 250401) at 10 µM was applied to the distal compartment of the MFC while maintaining a proximal-to-distal volume gradient.

### Inhibition of dynamin-dependent endocytosis

Dynasore (Sigma Aldrich, D7693) at 100nM was added to the distal compartment of the MFC while maintaining a proximal-to-distal volume gradient.

### Pull down assays

For the cell cultures pull downs experiments, 2 x 10^6^ COS7 cells were plated in 10 cm culture dishes. The following day, cells were transfected using calcium phosphate protocol with Flag-CRMP4/GFP-CRMP4/GFP-Delta-CRMP4 vector. The next day, cells were lysed, and proteins were extracted using lysis buffer containing PBS, 1% Triton-100X, and 1% protease and phosphatase inhibitors (Roche), followed by centrifugation and collection of the supernatant. At this point, immunoprecipitation preparation of the lysate was precleared with protein-A-agarose beads (Roche). Following overnight incubation with primary anti-flag antibody/anti DIC antibody, complexes were incubated with protein A agarose beads for 2 h at 4 °C and then precipitated and washed with PBS with 0.1% Triton X-100 (Sigma). Proteins were eluted by boiling in sample buffer and then subjected to western blot precipitation analysis with CRMP4/dynactin p150/Flag antibody(Sigma-Alderich F3165)/DIC. We used mouse IgG antibody as a control. For sciatic nerve pull downs, 12 sciatic nerves were pooled for each experiment. Here as well, the P90 sciatic nerve samples were first excised and homogenized in lysis buffer containing PBS and 1% protease and phosphatase inhibitors (Roche), followed by centrifugation and collection of the supernatant. Then we performed the pull-down assay using the technique described above. Under these conditions, pull downs were performed using DIC (Millipore MAB1618) and CRMP4 (Millipore AB5454) antibodies.

### Western blotting

Sciatic nerve axoplasm was isolated by excising and cutting sciatic nerves into short segments, followed by detergent-free buffer homogenized with PBS X1 protease and phosphatase inhibitors (Roche), followed by centrifugation and collection of the supernatant. Complete sciatic nerve extracts were achieved in the same manner with the exception of adding 1% Triton X-100. The protein concentration was determined using the Bio-Rad Protein Assay. Protein samples were denatured by boiling in SDS sample buffer and then electrophoresed in 8% polyacrylamide gels (SDS-PAGE). Proteins were transferred to a nitrocellulose membrane and then immunoblotted with appropriate primary antibodies: anti-CRMP4 - 1:2000 (Millipore AB5454) anti-DIC – 1:1000 (Millipore MAB1618) anti-p150 1:250 (BD Bioscience 611003) anti-Flag 1:4000 (Sigma-Alderich F3165) anti-Tubulin 1:10,000 (ab7291); and anti-tERK 1:10,000, diluted in 5% (w/v) Skim-milk (BD Difco) in TBS-T, followed by species-specific HRP-conjugated secondary antibodies (Jackson Laboratories) and visualized using a myECL imager (Thermo), according to the manufacturer’s instructions. ImageJ software was used for quantification.

### Sciatic nerve sectioning and immunostaining

Sciatic nerves of P90 mice were isolated and immediately fixed by using 4% PFA followed by 20% sucrose incubation. Then the samples were embedded by freezing in Tissue-Tek**®** OCT. Next, 10 µm lumbar sciatic nerve sections were prepared using Cryotome**™** FSE cryostat (Thermo-Fisher Scientific). Sections were rinsed in PBS, and then permeabilized with 0.1% Triton X-100, 5% Goat Serum (GS), 1 mg/mL Bovine Serum Albumin IgG, and protease free (BSA) in PBS. Primary antibodies against NFH (1:500) and CRMP4 (1:250) were diluted in blocking solution, 5% GS, 1 mg/mL BSA in PBS, and incubated overnight at 4°C. Samples were incubated with species-specific fluorescent secondary antibodies for 2 hours at room temperature. ProLong antifade medium (Molecular Probes) was added and the samples were covered with a #1.5, 18×18 mm cover slide.

### Immunostaining of cell cultures

Cultures were fixed in 4% paraformaldehyde and permeabilized with 0.1% Triton X-100, 5% GS, 1 mg/mL BSA in PBS. Samples were blocked for 1 hour with blocking medium containing 5% GS and 1 mg/mL BSA in PBS. Primary antibodies against Tau 1:100 (abcam, ab80579) NFH - 1:500 (Sigma-Aldrich N4142), PlexinA1 - 1:100 (Alomone lab, APR-081-F), CRMP4-1:100 (Millipore, AB5454), GAPDH 1:500 (abcam, ab9484), Tubulin 1:500 (abcam, ab7291), HB9 1:100 (IMGENEX, IMG-6549A) were diluted in blocking solution and incubated overnight at 4°C. Samples were incubated with species-specific fluorescent secondary antibodies for 2 hours at room temperature. For visualizing nuclei in myotubes, DAPI was used. In the MFC, after the staining protocol was completed, the MFC was peeled from the dish by gently pulling it from the proximal to the distal side.

### Proximity ligation assay

The proximity ligation assay (PLA) was used to visualize the co-localization of selected proteins; it was performed as previously described (Söderberg *et al*., 2008). Briefly, iPSc-derived MNs and murine-MN cultures were grown in the MFC on glass dishes for 18 and 5 DIV, respectively, and were then fixed in 4% PFA, at 4°C for 20 minutes. Subsequently, the samples were blocked and permeabilized with 5% Donkey Serum, 1% BSA, and 0.1% Triton X-100 in PBS for 1h and incubated with anti-CRMP4 and anti-DIC antibodies overnight at 4°C. Interactions (range ∼40nm) were detected by the proximity ligation assay Duolink kit (Sigma: PLA probe anti-mouse minus DUO92004, anti-rabbit plus DUO92002, and the detection kit Far Red). PLA was performed according to the manufacturer’s instructions. Coverslips were washed, mounted, and imaged by confocal microscopy. Half ligation samples were used as a negative control. The axonal PLA signal was quantified with ImageJ software using an axonal mask based on an endogenous mCherry/Rosa signal. The PLA puncta signal was quantified with the analyzed particle function of the software.

### Peptide design and insertion into MNs

The Dynein-CRMP4 blocking peptide design was based on previous findings by Amrimura et al., which pointed to 50 specific amino acid sequences responsible for CRMP2 binding to dynein (Arimura *et al*., 2009). Peptides were prepared by Alomone labs and GL Biochem. Peptide sequences are as follows:

| Name | Sequence | MW (Da) |
| --- | --- | --- |
| **Peptide-1** | TTMIIDHVVPEPE | **1480 Da** |
| **Peptide-2** | SSLTEAYEKWREWADGKS | **2143 Da** |
| **Peptide-3** | CCDYALHVDI | **1151 Da** |
| **Peptide-4** | THWNDSVKQ | **1114 Da** |

The peptides were inserted into axons by harsh pipetting. The recovered axons contain the peptide, as shown in Supplementary Figure 5. Tamra peptide was generously donated by Dr. Mike Fainzilber’s lab.

### Vectors

CRMP4 and ΔCRMP4 (containing deletion of the coding sequence 301-450bp) were sub-cloned in frame into the pLL3.7-GFP (Addgene) mammalian expression vector. Flag-CRMP4, used in the pull-down assays, was cloned into pCDNA3 vector (Invitrogen).

### Experimental design and statistical analysis

All statistical analyses were performed using GraphPad Prism v6.0. For two-group analysis, Student’s t-test or the Mann-Whitney test was used, as determined by a normality test. For multiple comparisons, Anova was used with the Tukey or Holm-Sidak post-hoc tests. All experiments include at least 3 biologically independent repeats, Significance was set at p<0.05.

## Abbreviations

ALS: Amyotrophic Lateral Sclerosis
BDNF: Brain-Derived Neurotrophic Factor
CBP: CRMP4 Binding Peptides
CNTF: Ciliary Neurotrophic Factor
CTX: Alexa Fluor 647-conjugated Cholera toxin subunit B
C9orf72: Chromosome 9 open reading frame 72
CRMPs: Collapsin Respond Mediator Proteins
DIV: Days In Vitro
Dyn-In: Ciliobrevin-D (Dynein Inhibitor)
GFP-ΔCRMP4: CRMP4 missing amino acids 100-150
GDNF: Glial-Derived Neurotrophic
IPSC: Induced Pluripotent Stem Cells
JNK: c-Jun N-terminal kinases
MFC: Microfluidic Chambers
MNs: Motor Neurons
NMJ: Neuromuscular Junction
NRP1: Neuropilin 1
PTEN: Phosphatase and tensin homolog
PLO: Poly D-L-ornithine
PLA: Proximity Ligation Assay
PDMS: Polydimethylsiloxane
PNB: Poor Neuorobasal medium (neurotrophin- and serum-free medium)
SOD1: Cu/Zn Superoxide Dismutase 1
Sema3A: Semaphorin3A
SN: Sciatic Nervep75^NTR^ - low-affinity nerve growth factor receptor

## Ethics approval and consent to participate

Animal experiments were performed under the supervision and approval of the Tel-Aviv University Committee for Animal Ethics. Human iPSC experiments were performed under the supervision and approval of the Tel-Aviv University Committee for Human Ethics.

## Availability of data and material

All data generated or analyzed during this study are included in this published article

## Author Contribution

Experiment were performed by RM, LA, RW and ET. Data were collected and analyzed by RM, LA. The study was designed, coordinated, and written by RM, LA, RW, ET, TP, SB, MB and EP. All authors read and approved the final manuscript.

## Acknowledgments

We thank Prof. Mike Fainzilber from the Weizmann Institute for the Tamra peptides. We also thank Prof. Eva Feldman and Prof. Stephen Goutman for obtaining the fibroblasts for the IPSC lines.

## Figure Legends

**Supplementary Figure 1:**
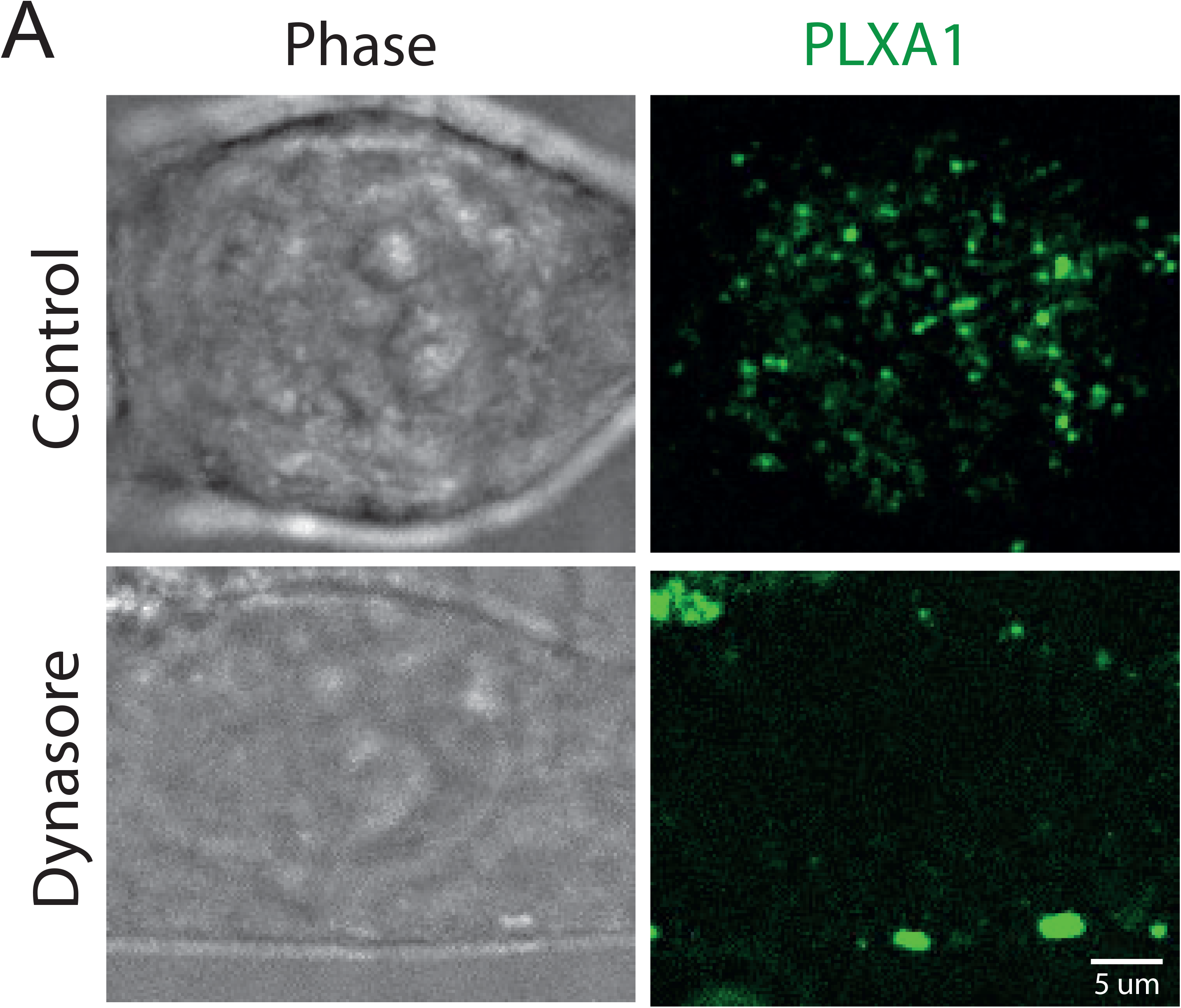
Dynasore blocks internalization of the Sema3A binding receptor. (A) Representative images of COS7 cells that were grown on glass dishes for 2DIV and then treated with FITC-PlexinA1 antibody with and without Dynasore. The Plexin A1-FITC antibody remained on the cell surface of cells that were treated with the dynamin-dependent endocytosis blocker compared with the untreated group.

**Supplementary Figure 2:**
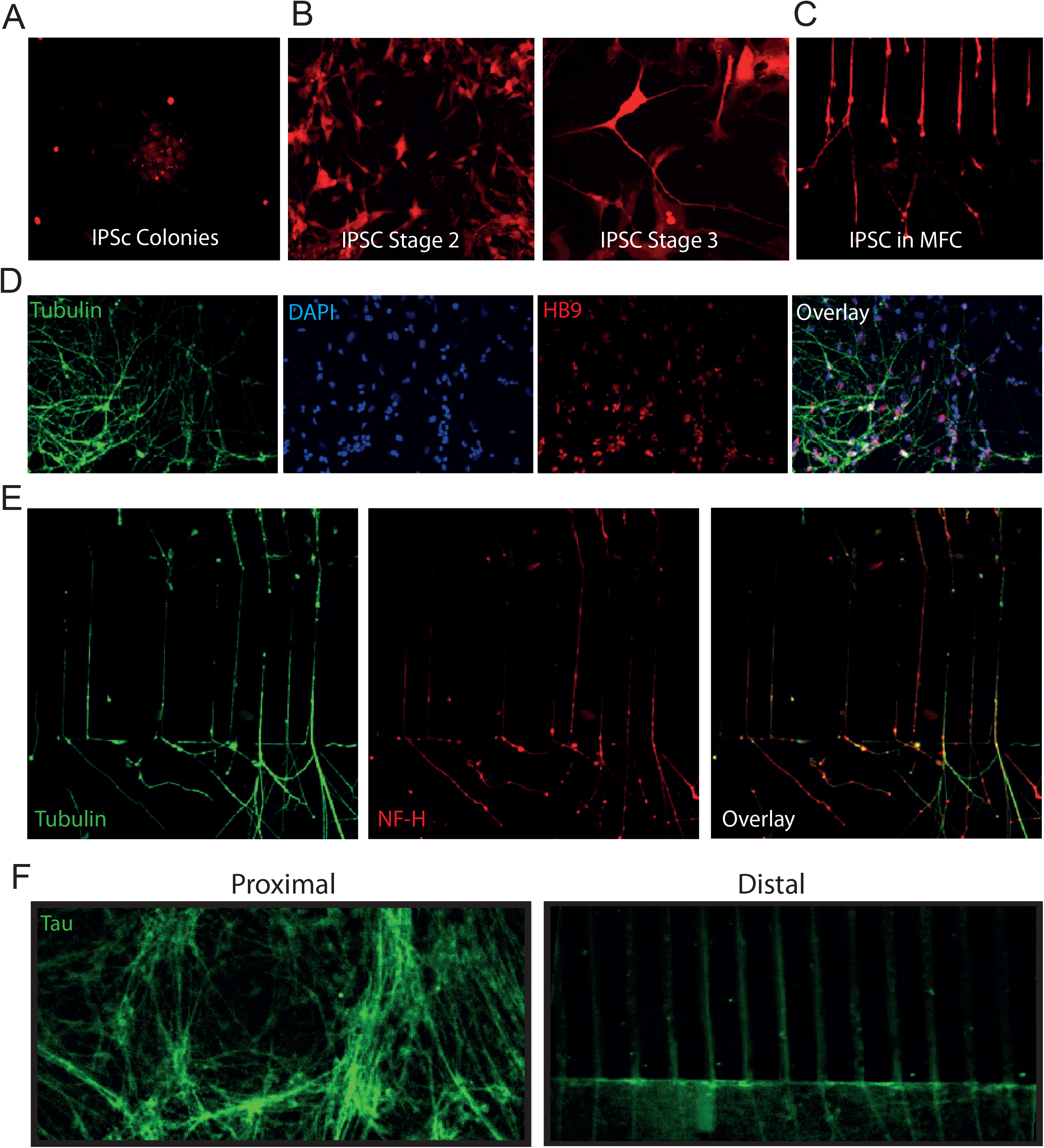
Human iPSC-derived motor neurons in a Microfluidic Device. (A) Representative image of C9orf72 mCherry-tagged iPSC colonies grown on Matrigel coating and treated with Nutristem media. The colonies were grown until they reach the right size for MN differentiation protocol. (B) Left panel - Representative image of differentiated C9orf72 mCherry-tagged iPSC colonies grown on laminin in our MFC treated with stage 2 differentiation media containing IMDM, F12, NEAA, B27, N2,PSA, LDN193189, SB431542, CHIR99021, All-Trans RA, and *SAG for another 6 DIV. Neuron-like morphology was achieved. Then the cells were treated with stage 3 media containing IMDM, F12, NEAA, B27, N2, PSA, Compound E, DAPT, db-cAMP, All-Trans RA, *SAG, and Ascorbic Acid until adult MN morphology and function were achieved (right panel). (C) Stage 3 iPSC-derived MNs reach out their axons into the microgrooves toward the proximal compartment after 18 DIV. (D) iPSC-derived MNs were fixed and stained at 18 DIV in the proximal compartment of the MFC for motor neuron specific markers. Green denotes tubulin, Blue denotes DAPI, and Red denotes HB9. Scale bar. (E) iPSC-derived MNs were fixed and stained at 18DIV in the distal compartment of the MFC for NFH and Tubulin. Scale bar. (F) Representative images for both proximal and distal compartments showing Tau-positive staining of our iPSC-derived MN cultures in both compartments.

**Supplementary Figure 3:**
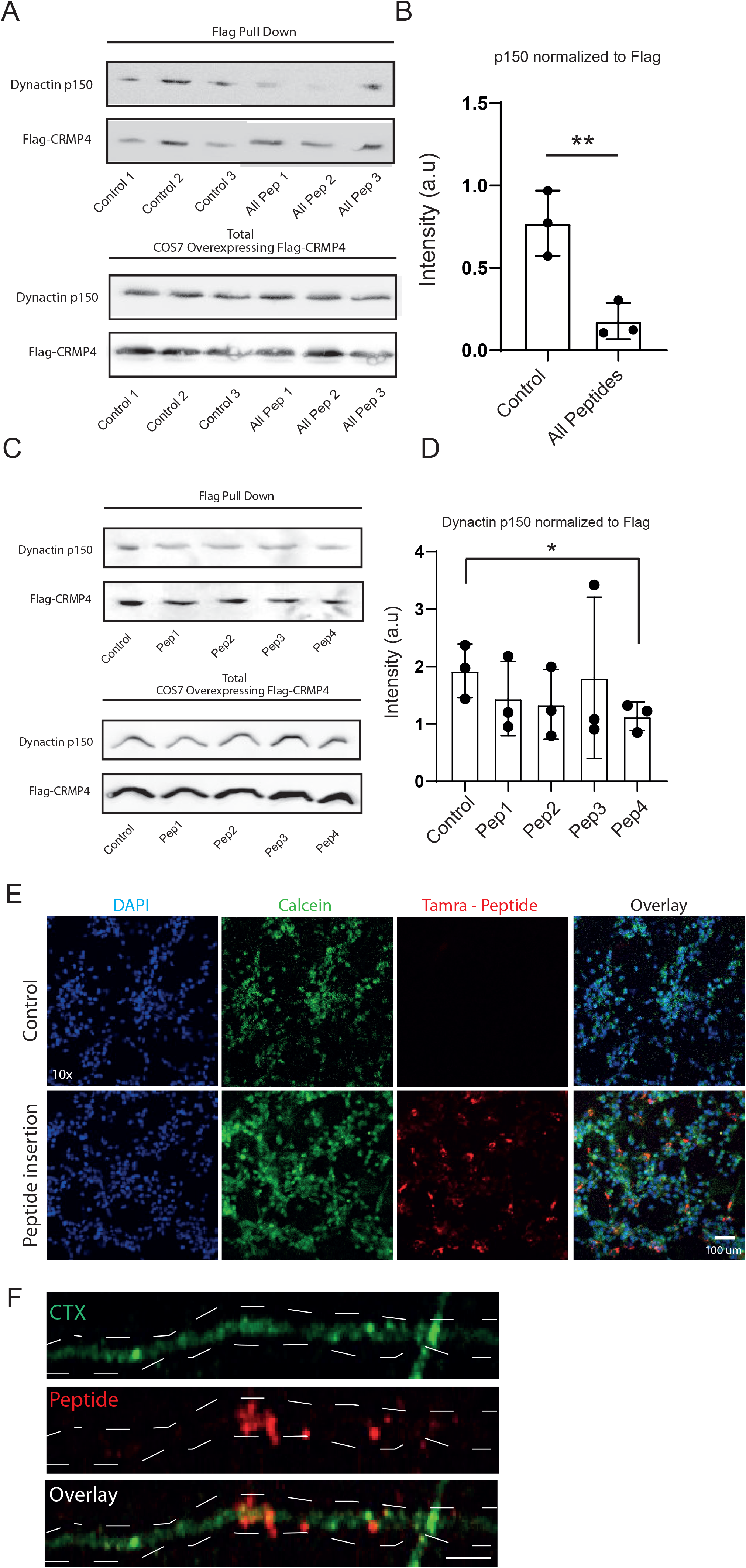
Peptide insertion into COS7 and MNs. (A) Upper panel - immunoprecipitation assay with anti-flag antibody followed by western blot analysis of dynactin (p150) showing a strong association of dynein complex with CRMP4 in COS7 overexpressing Flaf-CRMP4 protein. This binding can be interrupted with pre-incubation of the protein extract with a 10µm mixture of CBP1-4. Lower panel - Total protein western blot analysis indicates similar protein levels before the pull-down assay. (B) Quantification of the blot in A shows a significant decrease in Flag-CRMP4-dynactin binding when peptides were pre-incubated with the cell lysate compared with control. The dynactin intensity band normalized to the Flag-CRMP4 intensity band in each repeat (Student’s t-test, p=0.01, n=3). (C) Upper panel - Immunoprecipitation assay with anti-flag antibody followed by western blot analysis of dynactin (p150) showing a strong association of dynein complex with CRMP4 in COS7 overexpressing Flag-CRMP4 protein. This binding was tested by pre-incubation of the protein extract with 10µm of CBP1-4. Lower panel - Total protein western blot analysis indicates similar protein levels before the pull-down assay. (D) Quantification of the blot in C shows a mild but significant decrease in Flag-CRMP4-dynactin binding only when CBP4 was used, suggesting minimal effects on CRMP4-dynein complex binding for CBP1-3 in isolation. The dynactin intensity band normalized to the Flag-CRMP4 intensity band in each repeat (Student’s t-test, p=0.05, n=3). E) Representative images of MN culture after insertion of TAMRA peptides by harsh pipetting. The cells were stained with DAPI (Blue) and Calcein (Green). Red denotes TAMRA peptide. Scale bar: 100um. (F) Higher magnification images of MN axons in the distal compartment of an MFC after treatment with TAMRA peptide specifically in the axon compartment. Scale bar: 5um.

---

**Movie 1 – Dynein-dependent retrograde transport of Lysotracker and Mitotracker in MN axons.**

Time-lapse image series of lysosomal (green) and mitochondrial (red) trackers undergo dynein-dependent retrograde transport in MN axons. Scale bar: 20µm.

**Movie 2 – Dynein-dependent retrograde transport of Lysotracker and Mitotracker in MN axons is blocked by dynein inhibition.**

Time-lapse image series of lysosomal (green) and mitochondrial (red) trackers after applying a dynein inhibitor. The dynein-dependent retrograde transport in MN axons was blocked. Scale bar: 20µm.

**Movie 3 – Retrograde transport of biotin BDNF in axons of HB9::GFP MNs.**

Time-lapse image series of biotin BDNF, showing retrograde transport in HB9::GFP axons. Scale bar: 20µm.

**Movie 4 – Retrograde transport process of biotin BDNF in axons of HB9::GFP MN is blocked by applying Dynasore (an endocytosis inhibitor) in the distal compartment of an MFC device.**

Time-lapse image series of biotin BDNF after applying Dynosaore, an endocytosis inhibitor. The clathrin-dependent endocytosis event of BDNF was blocked; therefore, biotin BDNF did not undergo retrograde transport. Scale bar: 20µm.

## References

Alves, C. J. et al. (2015) ‘Gene expression profiling for human iPS-derived motor neurons from sporadic ALS patients reveals a strong association between mitochondrial functions and neurodegeneration’, Frontiers in Cellular Neuroscience, 9, p. 289. doi: 10.3389/fncel.2015.00289.

Arimura, N. et al. (2009) ‘CRMP-2 directly binds to cytoplasmic dynein and interferes with its activity’, Journal of Neurochemistry. Blackwell Publishing Ltd, 111(2), pp. 380–390. doi: 10.1111/j.1471-4159.2009.06317.x.

Balastik, M. et al. (2015) ‘Prolyl Isomerase Pin1 Regulates Axon Guidance by Stabilizing CRMP2A Selectively in Distal Axons’, Cell Reports. Cell Press, 13(4), pp. 812–828. doi: 10.1016/J.CELREP.2015.09.026.

Bamji, S. X. et al. (1998) ‘The p75 neurotrophin receptor mediates neuronal apoptosis and is essential for naturally occurring sympathetic neuron death.’, The Journal of cell biology, 140(4), pp. 911–23.

Ben-Zvi, A. et al. (2007) ‘Modulation of semaphorin3A activity by p75 neurotrophin receptor influences peripheral axon patterning.’, The Journal of neuroscience: the official journal of the Society for Neuroscience, 27(47), pp. 13000–11. doi: 10.1523/JNEUROSCI.3373-07.2007.

Ben-Zvi, A. et al. (2008) ‘The Semaphorin receptor PlexinA3 mediates neuronal apoptosis during dorsal root ganglia development.’, The Journal of neuroscience: the official journal of the Society for Neuroscience, 28(47), pp. 12427–32. doi: 10.1523/JNEUROSCI.3573-08.2008.

Bilsland, L. G. et al. (2010) ‘Deficits in axonal transport precede ALS symptoms in vivo.’, Proceedings of the National Academy of Sciences of the United States of America, 107(47), pp. 20523–8. doi: 10.1073/pnas.1006869107.

Birger, A. et al. (2018) ‘ALS-related human cortical and motor neurons survival is differentially affected by Sema3A’, Cell Death & Disease. Nature Publishing Group, 9(3), p. 256. doi: 10.1038/s41419-018-0294-6.

Blasco, H. et al. (2013) ‘A Rare Motor Neuron Deleterious Missense Mutation in the *DPYSL3* (*CRMP4*) Gene is Associated with ALS’, Human Mutation. John Wiley & Sons, Ltd, 34(7), pp. 953–960. doi: 10.1002/humu.22329.

Boillée, S., Vande Velde, C. and Cleveland, D. W. (2006) ‘ALS: A Disease of Motor Neurons and Their Nonneuronal Neighbors’, Neuron, 52(1), pp. 39–59. doi: 10.1016/j.neuron.2006.09.018.

Cagnetta, R. et al. (2018) ‘Rapid Cue-Specific Remodeling of the Nascent Axonal Proteome’, Neuron, 99(1), pp. 29–46.e4. doi: 10.1016/j.neuron.2018.06.004.

Cagnetta, R. et al. (2019) ‘Noncanonical Modulation of the eIF2 Pathway Controls an Increase in Local Translation during Neural Wiring’, Molecular Cell, 73(3), pp. 474–489.e5. doi: 10.1016/j.molcel.2018.11.013.

Campbell, D. S. and Holt, C. E. (2001) ‘Chemotropic responses of retinal growth cones mediated by rapid local protein synthesis and degradation.’, Neuron, 32(6), pp. 1013–26.

Castellani, V., Falk, J. and Rougon, G. (2004) ‘Semaphorin3A-induced receptor endocytosis during axon guidance responses is mediated by L1 CAM.’, Molecular and cellular neurosciences, 26(1), pp. 89–100. doi: 10.1016/j.mcn.2004.01.010.

Chadborn, N. H. et al. (2006) ‘PTEN couples Sema3A signalling to growth cone collapse.’, Journal of cell science, 119(Pt 5), pp. 951–7. doi: 10.1242/jcs.02801.

Charrier, E. et al. (2003) ‘Collapsin Response Mediator Proteins (CRMPs): Involvement in Nervous System Development and Adult Neurodegenerative Disorders’, Molecular Neurobiology. Humana Press, 28(1), pp. 51–64. doi: 10.1385/MN:28:1:51.

Cosker, K. E. and Segal, R. A. (2014) ‘Neuronal signaling through endocytosis.’, Cold Spring Harbor perspectives in biology, 6(2). doi: 10.1101/cshperspect.a020669.

Costa, C. J. and Willis, D. E. (2018) ‘To the end of the line: Axonal mRNA transport and local translation in health and neurodegenerative disease’, Developmental Neurobiology. John Wiley & Sons, Ltd, 78(3), pp. 209–220. doi: 10.1002/dneu.22555.

Dang, P., Smythe, E. and Furley, A. J. W. (2012) ‘TAG1 regulates the endocytic trafficking and signaling of the semaphorin3A receptor complex.’, The Journal of neuroscience: the official journal of the Society for Neuroscience, 32(30), pp. 10370– 82. doi: 10.1523/JNEUROSCI.5874-11.2012.

Deinhardt, K. et al. (2006) ‘Rab5 and Rab7 Control Endocytic Sorting along the Axonal Retrograde Transport Pathway’, Neuron, 52(2), pp. 293–305. doi: 10.1016/j.neuron.2006.08.018.

DeJesus-Hernandez, M. et al. (2011) ‘Expanded GGGGCC Hexanucleotide Repeat in Noncoding Region of C9ORF72 Causes Chromosome 9p-Linked FTD and ALS’, Neuron. Elsevier, 72(2), pp. 245–256. doi: 10.1016/J.NEURON.2011.09.011.

Devlin, A.-C. et al. (2015) ‘Human iPSC-derived motoneurons harbouring TARDBP or C9ORF72 ALS mutations are dysfunctional despite maintaining viability’, Nature Communications. Nature Publishing Group, 6(1), p. 5999. doi: 10.1038/ncomms6999.

Duplan, L. et al. (2010) ‘Collapsin response mediator protein 4a (CRMP4a) is upregulated in motoneurons of mutant SOD1 mice and can trigger motoneuron axonal degeneration and cell death.’, The Journal of neuroscience: the official journal of the Society for Neuroscience, 30(2), pp. 785–96. doi: 10.1523/JNEUROSCI.5411-09.2010.

Escudero, C. A. et al. (2019) ‘c-Jun N-terminal kinase (JNK)-dependent internalization and Rab5-dependent endocytic sorting mediate long-distance retrograde neuronal death induced by axonal BDNF-p75 signaling’, Scientific Reports. Nature Publishing Group, 9(1), p. 6070. doi: 10.1038/s41598-019-42420-6.

Fischer, L. R. et al. (2004) ‘Amyotrophic lateral sclerosis is a distal axonopathy: evidence in mice and man.’, Experimental neurology, 185(2), pp. 232–40.

Fournier, A. E. et al. (2000) ‘Semaphorin3a Enhances Endocytosis at Sites of Receptor–F-Actin Colocalization during Growth Cone Collapse’, The Journal of Cell Biology, 149(2), pp. 411–422. doi: 10.1083/jcb.149.2.411.

Frey, D. et al. (2000) ‘Early and selective loss of neuromuscular synapse subtypes with low sprouting competence in motoneuron diseases.’, The Journal of neuroscience: the official journal of the Society for Neuroscience, 20(7), pp. 2534– 42.

Fujimori, K. et al. (2018) ‘Modeling sporadic ALS in iPSC-derived motor neurons identifies a potential therapeutic agent’, Nature Medicine. Nature Publishing Group, 24(10), pp. 1579–1589. doi: 10.1038/s41591-018-0140-5.

Gershoni-Emek, N. et al. (2015) ‘Amyotrophic Lateral Sclerosis as a Spatiotemporal Mislocalization Disease: Location, Location, Location’, International Review of Cell and Molecular Biology. Academic Press, 315, pp. 23–71. doi: 10.1016/BS.IRCMB.2014.11.003.

Gibbs, K. L. et al. (2018) ‘Inhibiting p38 MAPK alpha rescues axonal retrograde transport defects in a mouse model of ALS’, Cell Death & Disease. Nature Publishing Group, 9(6), p. 596. doi: 10.1038/s41419-018-0624-8.

Good, P. F. et al. (2004) ‘A role for semaphorin 3A signaling in the degeneration of hippocampal neurons during Alzheimer’s disease.’, Journal of neurochemistry, 91(3), pp. 716–36. doi: 10.1111/j.1471-4159.2004.02766.x.

Goshima, Y. et al. (1997) ‘A novel action of collapsin: collapsin-1 increases antero- and retrograde axoplasmic transport independently of growth cone collapse.’, Journal of neurobiology, 33(3), pp. 316–28.

Haramati, S. et al. (2010) ‘miRNA malfunction causes spinal motor neuron disease.’, Proceedings of the National Academy of Sciences of the United States of America, 107(29), pp. 13111–6. doi: 10.1073/pnas.1006151107.

Harrington, A. W. and Ginty, D. D. (2013) ‘Long-distance retrograde neurotrophic factor signalling in neurons.’, Nature reviews. Neuroscience. Nature Publishing Group, a division of Macmillan Publishers Limited. All Rights Reserved., 14(3), pp. 177–87. doi: 10.1038/nrn3253.

Howard, J., Hudspeth, A. J. and Vale, R. D. (1989) ‘Movement of microtubules by single kinesin molecules’, Nature. Nature Publishing Group, 342(6246), pp. 154–158. doi: 10.1038/342154a0.

Ibáñez, C. F. (2007) ‘Message in a bottle: long-range retrograde signaling in the nervous system’, Trends in Cell Biology. Elsevier Current Trends, 17(11), pp. 519–528. doi: 10.1016/J.TCB.2007.09.003.

Ionescu, A. et al. (2016) ‘Compartmental microfluidic system for studying muscle– neuron communication and neuromuscular junction maintenance’, European Journal of Cell Biology, 95(2), pp. 69–88. doi: 10.1016/j.ejcb.2015.11.004.

Ionescu, A. et al. (2019) ‘Targeting the Sigma-1 Receptor via Pridopidine Ameliorates Central Features of ALS Pathology in a SOD1G93A Model’, Cell Death & Disease. Nature Publishing Group, 10(3), p. 210. doi: 10.1038/s41419-019-1451-2.

Kenchappa, R. S. et al. (2010) ‘p75 neurotrophin receptor-mediated apoptosis in sympathetic neurons involves a biphasic activation of JNK and up-regulation of tumor necrosis factor-alpha-converting enzyme/ADAM17.’, The Journal of biological chemistry, 285(26), pp. 20358–68. doi: 10.1074/jbc.M109.082834.

Körner, S. et al. (2016) ‘The Axon Guidance Protein Semaphorin 3A Is Increased in the Motor Cortex of Patients With Amyotrophic Lateral Sclerosis.’, Journal of neuropathology and experimental neurology. The Oxford University Press, p. nlw003. doi: 10.1093/jnen/nlw003.

LaMonte, B. H. et al. (2002) ‘Disruption of dynein/dynactin inhibits axonal transport in motor neurons causing late-onset progressive degeneration.’, Neuron, 34(5), pp. 715–27.

Lee, S. and Huang, E. J. (2017) ‘Modeling ALS and FTD with iPSC-derived neurons’, Brain Research. Elsevier, 1656, pp. 88–97. doi: 10.1016/J.BRAINRES.2015.10.003.

Luo, Y., Raible, D. and Raper, J. A. (1993) ‘Collapsin: a protein in brain that induces the collapse and paralysis of neuronal growth cones.’, Cell, 75(2), pp. 217–27.

Maimon, R. et al. (2018) ‘miR126-5p Downregulation Facilitates Axon Degeneration and NMJ Disruption via a Non-Cell-Autonomous Mechanism in ALS.’, The Journal of neuroscience: the official journal of the Society for Neuroscience. Society for Neuroscience, 38(24), pp. 5478–5494. doi: 10.1523/JNEUROSCI.3037-17.2018.

Maimon, R. and Perlson, E. (2019) ‘Muscle secretion of toxic factors, regulated by miR126-5p, facilitates motor neuron degeneration in amyotrophic lateral sclerosis’, Neural Regeneration Research, 14(6), p. 969. doi: 10.4103/1673-5374.250571.

Manns, R. P. C. et al. (2012) ‘Differing semaphorin 3A concentrations trigger distinct signaling mechanisms in growth cone collapse.’, The Journal of neuroscience: the official journal of the Society for Neuroscience. Europe PMC Funders, 32(25), pp. 8554–9. doi: 10.1523/JNEUROSCI.5964-11.2012.

Millecamps, S. and Julien, J.-P. (2013) ‘Axonal transport deficits and neurodegenerative diseases’, Nature Reviews Neuroscience. Nature Publishing Group, 14(3), pp. 161–176. doi: 10.1038/nrn3380.

Molofsky, A. V et al. (2014) ‘Astrocyte-encoded positional cues maintain sensorimotor circuit integrity.’, Nature, 509(7499), pp. 189–94. doi: 10.1038/nature13161.

Moloney, E. B. et al. (2017) ‘Expression of a Mutant SEMA3A Protein with Diminished Signalling Capacity Does Not Alter ALS-Related Motor Decline, or Confer Changes in NMJ Plasticity after BotoxA-Induced Paralysis of Male Gastrocnemic Muscle’, PLOS ONE. Edited by B. Key. Public Library of Science, 12(1), p. e0170314. doi: 10.1371/journal.pone.0170314.

Moloney, E. B., de Winter, F. and Verhaagen, J. (2014) ‘ALS as a distal axonopathy: molecular mechanisms affecting neuromuscular junction stability in the presymptomatic stages of the disease.’, Frontiers in neuroscience, 8, p. 252. doi: 10.3389/fnins.2014.00252.

Münch, C. et al. (2004) ‘Point mutations of the p150 subunit of dynactin (DCTN1) gene in ALS.’, Neurology. Wolters Kluwer Health, Inc. on behalf of the American Academy of Neurology, 63(4), pp. 724–6. doi: 10.1212/01.wnl.0000134608.83927.b1.

Nagai, J. et al. (2015) ‘Crmp4 deletion promotes recovery from spinal cord injury by neuroprotection and limited scar formation’, Scientific Reports. Nature Publishing Group, 5(1), p. 8269. doi: 10.1038/srep08269.

Nagai, J., Baba, R. and Ohshima, T. (2017) ‘CRMPs Function in Neurons and Glial Cells: Potential Therapeutic Targets for Neurodegenerative Diseases and CNS Injury’, Molecular Neurobiology, 54(6), pp. 4243–4256. doi: 10.1007/s12035-016-0005-1.

Nakamura, F., Kalb, R. G. and Strittmatter, S. M. (2000) ‘Molecular basis of semaphorin-mediated axon guidance.’, Journal of neurobiology, 44(2), pp. 219–29.

Nicolas, A. et al. (2018) ‘Genome-wide Analyses Identify KIF5A as a Novel ALS Gene’, Neuron. Cell Press, 97(6), pp. 1268–1283.e6. doi: 10.1016/J.NEURON.2018.02.027.

Olenick, M. A., Dominguez, R. and Holzbaur, E. L. F. (2019) ‘Dynein activator Hook1 is required for trafficking of BDNF-signaling endosomes in neurons’, The Journal of Cell Biology, 218(1), pp. 220–233. doi: 10.1083/jcb.201805016.

Paschal, B. M. and Vallee, R. B. (1987) ‘Retrograde transport by the microtubule-associated protein MAP 1C’, Nature. Nature Publishing Group, 330(6144), pp. 181– 183. doi: 10.1038/330181a0.

Pathak, A. et al. (2018) ‘Retrograde Degenerative Signaling Mediated by the p75 Neurotrophin Receptor Requires p150Glued Deacetylation by Axonal HDAC1’, Developmental Cell. Cell Press, 46(3), pp. 376–387.e7. doi: 10.1016/J.DEVCEL.2018.07.001.

Perlson, E. et al. (2009) ‘A switch in retrograde signaling from survival to stress in rapid-onset neurodegeneration.’, The Journal of neuroscience: the official journal of the Society for Neuroscience, 29(31), pp. 9903–17. doi: 10.1523/JNEUROSCI.0813-09.2009.

Perlson, E. et al. (2010) ‘Retrograde axonal transport: pathways to cell death?’, Trends in neurosciences, 33(7), pp. 335–44. doi: 10.1016/j.tins.2010.03.006.

Peters, O. M., Ghasemi, M. and Brown, R. H. (2015) ‘Emerging mechanisms of molecular pathology in ALS.’, The Journal of clinical investigation, 125(5), pp. 1767– 79. doi: 10.1172/JCI71601.

Ponnusamy, R. et al. (2014) ‘Crystal Structure of Human Crmp-4: Correction of Intensities for Lattice-Translocation Disorder’, Acta Crystallogr.,Sect.D, 70, p. 1680. doi: 10.2210/PDB4CNT/PDB.

Rahajeng, J. et al. (2010) ‘Collapsin Response Mediator Protein-2 (Crmp2) Regulates Trafficking by Linking Endocytic Regulatory Proteins to Dynein Motors’, Journal of Biological Chemistry, 285(42), pp. 31918–31922. doi: 10.1074/jbc.C110.166066.

Renton, A. E. et al. (2011) ‘A Hexanucleotide Repeat Expansion in C9ORF72 Is the Cause of Chromosome 9p21-Linked ALS-FTD’, Neuron, 72(2), pp. 257–268. doi: 10.1016/j.neuron.2011.09.010.

Rosen, D. R. et al. (1993) ‘Mutations in Cu/Zn superoxide dismutase gene are associated with familial amyotrophic lateral sclerosis’, Nature, 362(6415), pp. 59–62. doi: 10.1038/362059a0.

Rotem, N. et al. (2017) ‘ALS Along the Axons – Expression of Coding and Noncoding RNA Differs in Axons of ALS models’, Scientific Reports. Nature Publishing Group, 7, p. 44500. doi: 10.1038/srep44500.

Sances, S. et al. (2016) ‘Modeling ALS with motor neurons derived from human induced pluripotent stem cells’, Nature Neuroscience. Nature Publishing Group, 19(4), pp. 542–553. doi: 10.1038/nn.4273.

Sasaki, Y. et al. (2002) ‘Fyn and Cdk5 Mediate Semaphorin-3A Signaling, Which Is Involved in Regulation of Dendrite Orientation in Cerebral Cortex’, Neuron. Cell Press, 35(5), pp. 907–920. doi: 10.1016/S0896-6273(02)00857-7.

Schmidt, E. F. and Strittmatter, S. M. (2007) ‘The CRMP Family of Proteins and Their Role in Sema3A Signaling’, in Semaphorins: Receptor and Intracellular Signaling Mechanisms. New York, NY: Springer New York, pp. 1–11. doi: 10.1007/978-0-387-70956-7_1.

Shi, Y. et al. (2018) ‘Haploinsufficiency leads to neurodegeneration in C9ORF72 ALS/FTD human induced motor neurons’, Nature Medicine. Nature Publishing Group, 24(3), pp. 313–325. doi: 10.1038/nm.4490.

Singh, K. K. et al. (2008) ‘Developmental axon pruning mediated by BDNF-p75NTR-dependent axon degeneration.’, Nature neuroscience, 11(6), pp. 649–58. doi: 10.1038/nn.2114.

Söderberg, O. et al. (2008) ‘Characterizing proteins and their interactions in cells and tissues using the in situ proximity ligation assay’, Methods. Academic Press, 45(3), pp. 227–232. doi: 10.1016/J.YMETH.2008.06.014.

Song, W. et al. (2010) ‘ProNGF induces PTEN via p75NTR to suppress Trk-mediated survival signaling in brain neurons.’, The Journal of neuroscience: the official journal of the Society for Neuroscience, 30(46), pp. 15608–15. doi: 10.1523/JNEUROSCI.2581-10.2010.

Steinberg, K. M. et al. (2015) ‘Exome sequencing of case-unaffected-parents trios reveals recessive and de novo genetic variants in sporadic ALS’, Scientific Reports. Nature Publishing Group, 5(1), p. 9124. doi: 10.1038/srep09124.

Tank, E. M. et al. (2018) ‘Abnormal RNA stability in amyotrophic lateral sclerosis’, Nature Communications. Nature Publishing Group, 9(1), p. 2845. doi: 10.1038/s41467-018-05049-z.

Terenzio, M., Schiavo, G. and Fainzilber, M. (2017) ‘Compartmentalized Signaling in Neurons: From Cell Biology to Neuroscience’, Neuron. Cell Press, 96(3), pp. 667– 679. doi: 10.1016/J.NEURON.2017.10.015.

Valdez, G. et al. (2012) ‘Shared resistance to aging and ALS in neuromuscular junctions of specific muscles.’, PloS one. Public Library of Science, 7(4), p. e34640. doi: 10.1371/journal.pone.0034640.

Venkova, K. et al. (2014) ‘Semaphorin 3A signaling through neuropilin-1 is an early trigger for distal axonopathy in the SOD1G93A mouse model of amyotrophic lateral sclerosis.’, Journal of neuropathology and experimental neurology. The Oxford University Press, 73(7), pp. 702–13. doi: 10.1097/NEN.0000000000000086.

De Vos, K. J. and Hafezparast, M. (2017) ‘Neurobiology of axonal transport defects in motor neuron diseases: Opportunities for translational research?’, Neurobiology of Disease. Academic Press, 105, pp. 283–299. doi: 10.1016/J.NBD.2017.02.004.

Wehner, A. B. et al. (2016) ‘Semaphorin 3A is a retrograde cell death signal in developing sympathetic neurons.’, Development (Cambridge, England), 143(9), pp. 1560–70. doi: 10.1242/dev.134627.

Wen, X. et al. (2014) ‘Antisense Proline-Arginine RAN Dipeptides Linked to C9ORF72-ALS/FTD Form Toxic Nuclear Aggregates that Initiate In Vitro and In Vivo Neuronal Death’, Neuron, 84(6), pp. 1213–1225. doi: 10.1016/j.neuron.2014.12.010.

De Winter, F. et al. (2006) ‘The expression of the chemorepellent Semaphorin 3A is selectively induced in terminal Schwann cells of a subset of neuromuscular synapses that display limited anatomical plasticity and enhanced vulnerability in motor neuron disease.’, Molecular and cellular neurosciences, 32(1–2), pp. 102–17. doi: 10.1016/j.mcn.2006.03.002.

de Wit, J. et al. (2006) ‘Vesicular trafficking of semaphorin 3A is activity-dependent and differs between axons and dendrites.’, Traffic (Copenhagen, Denmark), 7(8), pp. 1060–77. doi: 10.1111/j.1600-0854.2006.00442.x.

Wu, K. Y. et al. (2005) ‘Local translation of RhoA regulates growth cone collapse.’, Nature. NIH Public Access, 436(7053), pp. 1020–1024. doi: 10.1038/nature03885.

Yamashita, N. et al. (2014) ‘Plexin-A4-dependent retrograde semaphorin 3A signalling regulates the dendritic localization of GluA2-containing AMPA receptors.’, Nature communications, 5, p. 3424. doi: 10.1038/ncomms4424.

Yamashita, N. and Goshima, Y. (2012) ‘Collapsin Response Mediator Proteins Regulate Neuronal Development and Plasticity by Switching Their Phosphorylation Status’, Molecular Neurobiology, 45(2), pp. 234–246. doi: 10.1007/s12035-012-8242- 4.

Zahavi, E. E. et al. (2015) ‘A compartmentalized microfluidic neuromuscular co-culture system reveals spatial aspects of GDNF functions.’, Journal of cell science, 128(6), pp. 1241–52. doi: 10.1242/jcs.167544.

Zahavi, E. E., Maimon, R. and Perlson, E. (2017) ‘Spatial-specific functions in retrograde neuronal signalling’, Traffic. John Wiley & Sons A/S. doi: 10.1111/tra.12487.

